# Nectar chemistry reflects pollination strategies in alpine plant communities

**DOI:** 10.64898/2026.07.24.740324

**Authors:** Roberto Rebollo, Yan Yang, James W. Sims, Lea Stern, Mark C. Mescher, Consuelo M. De Moraes, Pengjuan Zu

## Abstract

Nectar is a primary floral reward, which can also deter non-specialist pollinators; yet, the role of complex nectar composition in plant–pollinator interactions and its relationship to other floral traits remain understudied, particularly in natural communities. We profiled the nectar metabolome of 43 co-occurring species in an alpine grassland to evaluate relationships among nectar chemistry, phylogeny, floral display traits and pollinator visitation. Field observations revealed a dominant bee vs fly visitation gradient. Several aggregated nectar traits tracked this gradient independently of plant phylogeny, indicating functional convergence: specifically, bee visitation was associated with higher nectar sucrose proportion, greater compound richness, and more divergent profiles of uncommon sugars. Nectar traits exhibited variable relationships to floral display traits, suggesting partial inter-trait phenotypic integration. Together, these findings indicate that detailed features of nectar composition, including phytochemical diversity, play a functional role in floral phenotypes, with previously unappreciated implications for plant–pollinator community ecology.

## Introduction

The adaptive strategies by which plants selectively attract effective pollinators while filtering out less effective ones are a central topic in pollination ecology, with direct implications for interaction networks and plant reproduction. Nectar is the most widespread floral reward offered to pollinators (Ballarin *et al*., 2024) and plays a central role in mediating these interactions: sugar compounds provide the energetic reward that broadly attracts visitors, while a richer blend of primary and secondary metabolites selectively filters which visitors are recruited or deterred (Nepi, 2014; Nicolson, 2022; Stevenson *et al*., 2017; Tiedge & Lohaus, 2017). For example, secondary compounds such as alkaloids can deter non-specialist or robbing visitors while reinforcing fidelity in effective pollinators (Leonhardt *et al*., 2024; Stevenson, 2020), whereas amino acids and non-standard sugars can be key in recruiting specific pollinators (Gegear *et al*., 2007; Nicolson & Thornburg, 2007). Because co-occurring species share potential pollinators — and may therefore experience reproductive competition and facilitation that influence community assembly and floral trait evolution (E-Vojtkó *et al*., 2020; Sargent & Ackerly, 2008) — reward chemistry is best studied in a community context (Barberis *et al*., 2023). Yet cross-species comparison of nectar chemistry has rarely been extended beyond simple sugar ratios (Abrahamczyk *et al*., 2017; Chalcoff *et al*., 2017; Liu *et al*., 2024; Perret *et al*., 2001). Moreover, despite increasing availability of sophisticated analytic tools (Roy *et al*., 2017), metabolomic studies have remained restricted to specific taxonomic groups (MacNeill *et al*., 2025) or species not found in the same community (Palmer-Young *et al*., 2019). These limitations leave a critical gap in our understanding of how the chemistry of floral rewards influences plant–pollinator interactions in natural communities.

Alpine systems provide an intriguing system for evaluating the relationship between floral traits and pollination strategies, as they typically exhibit high plant species diversity within a geographically compact area (Väre *et al*., 2003) and are dominated by two ecologically contrasting groups of insect pollinators: bees (clade Anthophila) and flies (order Diptera), which have distinct perceptual systems (Arnold *et al*., 2009), and flower-visiting behaviour (Apland & Koski, 2025; Herrera, 1987, 1990). These groups also differ in their degree of trophic specialisation, with bees being obligate flower visitors that form specialized relationships with many different plant species (Leonhardt & Blüthgen, 2012; Polidori *et al*., 2025; Wood *et al*., 2021), while flies are largely opportunistic and generalised flower visitors (Kearns, 1992; Larson *et al*., 2001; Lefebvre *et al*., 2014; Oldroyd, 1964; Tiusanen *et al*., 2016). Correspondingly, pollination by each of these insect groups is linked to different suites of floral traits, including flower shapes, colours, scents, and other features (Dellinger, 2020; Faegri & Van Der Pijl, 1979; Kantsa *et al*., 2018; Rosas-Guerrero *et al*., 2014). It seems likely that similar variation extends to the chemical composition of nectar, but such patterns remain poorly characterised.

Indeed, relatively little is known about the chemical features of nectar associated with pollination by bees or flies, apart from the proportion of sucrose, which is widely observed to be high in bee-associated nectar (Abrahamczyk *et al*., 2017; Baker & Baker, 1983; Liu *et al*., 2024; Vandelook *et al*., 2019). Another key nectar feature that could influence associations with these pollinator groups is phytochemical diversity. In the broader context of plant-insect interactions, higher metabolite diversity in vegetative plant tissues has been implicated in the mediation of simultaneous biotic interactions, particularly the deterrence of non-specialised herbivores (Berenbaum & Zangerl, 1996; reviewed in Kessler & Kalske, 2018). This pattern has received empirical support at multiple ecological scales, including natural and experimental communities (Abdala-Roberts & Moreira, 2024; Hauri *et al*., 2021; Salazar *et al*., 2016, 2018). Metabolite diversity in floral rewards may play an analogous role in filtering out non-specialist pollinators: for example, richer, more chemically divergent nectar profiles may simultaneously enhance attractiveness for effective pollinators while deterring opportunistic or ineffective visitors.

Supporting this hypothesis, several uncommon nectar components, such as nectar pigments, volatile organic compounds, and non-protein amino acids, have been linked to interactions with particular pollinator groups (often specialists) (Hansen *et al*., 2007; Stevenson *et al*., 2017; Wong *et al*., 2023). Uncommon nectar sugars (i.e., those other than sucrose, glucose, and fructose) are of particular interest in this regard, as they can have toxic or deterrent effects on some pollinators (Allsopp *et al*., 1998; Barker, 1977; Barker & Lehner, 1974; von Frisch, 1971; Roy *et al*., 2017; Sols *et al*., 1960). Based on these patterns, we predicted that alpine plants with higher nectar sucrose proportion, compound-rich profiles and higher multivariate dispersion (more divergent composition) would exhibit bee-dominated pollination strategies.

To explore whether and how nectar traits match plant pollination strategies while controlling for plant phylogenetic relatedness, we first described the nectar composition of a comprehensive set of 43 species (285 individual plants across 38 genera and 19 families) from two alpine grassland sites on a single mountain in eastern Switzerland. As discussed above, we focused on several *aggregated* nectar traits as potential indicators of the bee/fly pollination strategy: sucrose proportion, chemical richness, and multivariate dispersion, with richness and dispersion variables calculated separately for the overall metabolome and uncommon sugars. We evaluated whether these nectar traits correlated with field visitation patterns, whether they differed among floral display categories (morphology and colour), and whether display categories themselves tracked insect visitation, thereby indicating concerted pollination strategies.

## Material and methods

### Field sites

Our field sites comprised two grasslands on the Calanda mountain, Canton Grisons, Switzerland: one subalpine (1400 m a.s.l. 46°52’09.1”N, 9°29’25.0”E) and an alpine site (2000 m a.s.l., 46°53’15.9”N, 9°29’21.3”E), with 2.05 km of horizontal separation. Within each site, sampling took place within a 50 m radius in each location. Both sites have calcareous soils and are on southeast-facing slopes of similar inclination. The average temperature at these elevations is 7.0 °C and 4.4 °C, respectively, with mean annual precipitation of 1169 mm and 1355 mm (Alexander *et al*., 2015).

To comprehensively cover the diversity of insect-pollinated species in these species-rich communities, we selected all abundant plant species on which nectar collection was feasible. This resulted in a total of 43 species, belonging to 38 genera and 19 families (**Table S1**). Of these, 13 species were sampled at both locations. We note that some abundant species, including *Crocus albiflorus, Helianthemum alpestre, Helianthemum nummularium*, and *Plantago media*, were excluded from the study as they were not observed to produce nectar.

### Nectar collection

Nectar collection was conducted throughout the flowering season from May to August in 2023. To sample flower nectar, we first covered the flowers or inflorescences with fine mesh bags for approximately 24 hours prior to collection, to prevent depletion by visiting insects. Nectar was then collected from freshly open flowers using glass microcapillary tubes of various volumes (5, 1, and 0.5 µl) depending on the size of the flowers and the amount of nectar they produced. We recorded nectar volumes and the number of flowers we collected from. For 6 species with extremely viscous or crystallised nectar (*Alchemilla conjuncta, Alchemilla xanthochlora, Carum carvi, Dryas octopetala, Galium pumilum* and *Saxifraga paniculata*), we washed the nectar with a measured volume of double-distilled water using a micropipette, following the method from Power *et al*. (2018). For both collection methods, we pooled nectar from flowers of the same plant individual and transferred the samples into 1.5 mL Eppendorf vials (Eppendorf AG, Hamburg, Germany) pre-filled with 80 µL analysis-grade ethanol to mitigate degradation (this ethanol was evaporated at the beginning of the analysis protocol). Samples were kept in a cool box during fieldwork and transported to a -80°C freezer for storage until analysis. All nectar samples were collected on fully or mostly sunny days to ensure typical nectar production. We sampled on average 6.2 samples per species (for sample size per species, see **Figure 1**). For three species (*Gymnadenia nigra, Carduus defloratus*, and *Pilosella officinarum*), only two samples were obtained due to low nectar volume and the labour-intensive handling of small florets.

**Figure 1:**
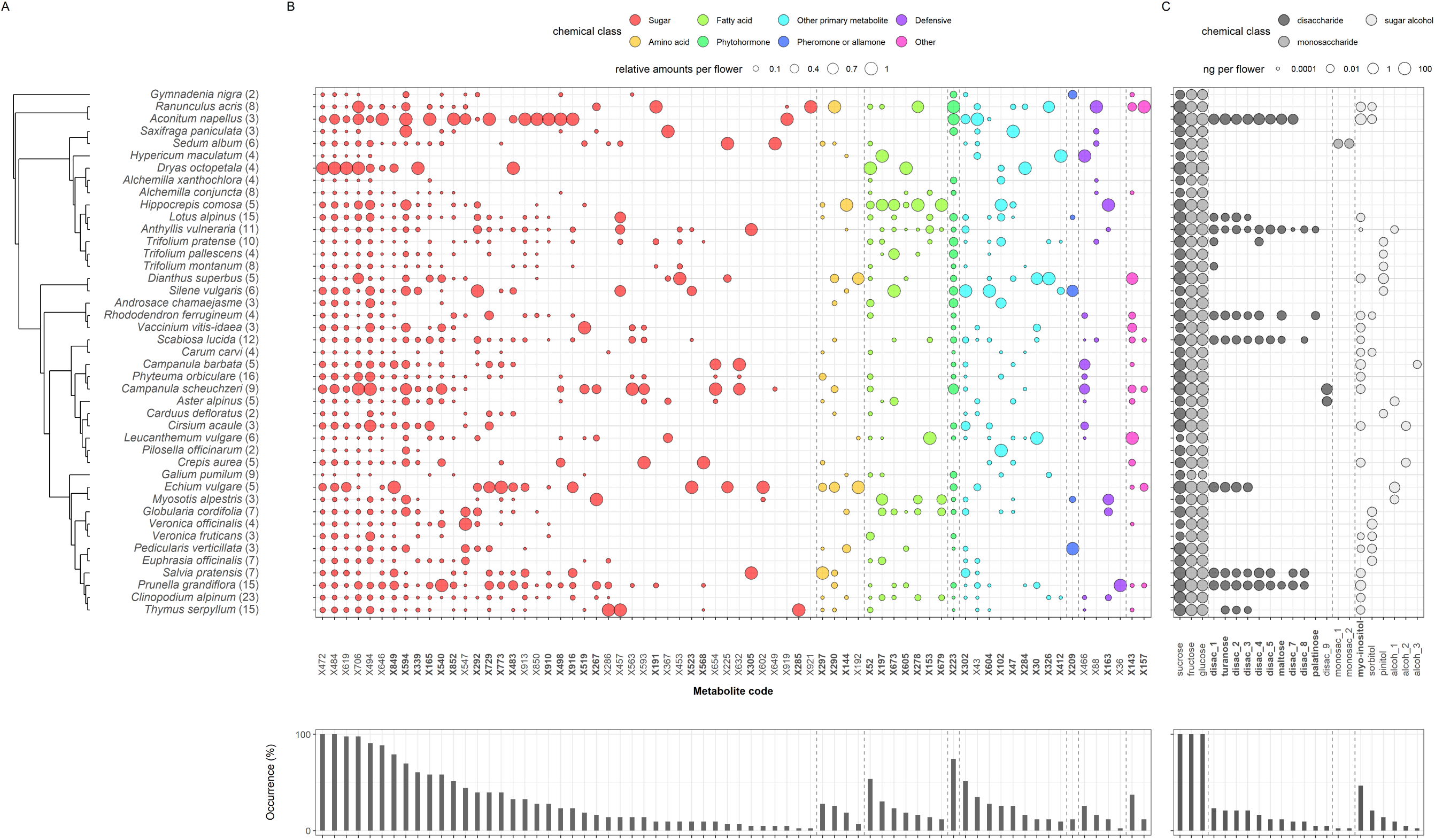
Nectar metabolomic profile of species in this study. **A**. Phylogenetic tree of species in the study, with their sample size in parentheses. **B**. untargeted MS-based analysis. **C**. FID quantification for sugars and sugar-related compounds. Metabolite codes in bold mark those with a significant phylogenetic signal (FDR-corrected *p* < 0.05, Table S3). The frequency of each metabolite, defined as the proportion of plant species in which it was found, is displayed at the bottom.

### Nectar metabolomic analyses

Nectar was analysed using an untargeted GC-MS metabolomics protocol described by Lisec *et al*. (2006) that involved compound derivatisation. For each nectar sample, we ran one split (1:20) injection to capture predominant sugars while avoiding over-saturation, and one splitless injection to better capture low-abundance metabolites. Briefly, splitless runs were analysed using Mzmine (4.3.3) (Heuckeroth *et al*., 2024), followed by manual verification of compounds identified to at least the chemical class level. Meanwhile, split runs were analysed using the MassHunter software package (version B.07.00, Agilent, USA) to identify and quantify the predominant sugars. For method and instrument details on the metabolomic analysis, see **Methods S1**.

### Calculation of aggregated nectar traits

We calculated the five aggregated nectar traits, which we predicted to be indicative of the plant pollination strategy (as previously discussed), at the sample level. Calculations were performed with R (R Core Team, 2021), version 4.4.2 (2024-10-31). Nectar sucrose proportion was calculated as the sucrose amount relative to the total main sugars (sucrose, fructose and glucose). Metabolite richness was the number of metabolites detected, and was done separately for overall metabolites and uncommon sugars, derived from splitless and split data, respectively.

The multivariate dispersion in chemical profiles was also calculated separately for overall metabolites and uncommon sugars. For each of these datasets, we first obtained a Jaccard distance matrix (using vegdist, from the vegan package; Oksanen *et al*., 2025), based on the presence/absence of individual metabolites in each sample. We then calculated the distance of all samples from the global median in this multi-dimensional trait space using the betadisper function (Oksanen *et al*., 2025), which served as our dispersion metric.

### Insect pollinator visitation

We monitored insect visitation during the flowering season (approximately from late April to August) in 2022 and 2023 using a combination of random walks and species-specific observations. Random walks were conducted across the sampling area under sunny weather, during which we recorded pollinator taxa and the plant species they visited using a notebook or camera. In total, we dedicated approximately 74 person-hours to random walks and an additional 11.25 person-hours to plant species with few recorded interactions.

In total, we considered 975 visitation events for the focal plant species. Only plant species with 5 or more observations were included in analyses evaluating visitation patterns (35 out of 43 species, median observations per species = 25, **Table S2**).

The observed taxonomical orders were: Diptera, Lepidoptera, Coleoptera and Hymenoptera. Hemipterans contributed to only three visits and were excluded from further analysis. From these orders, we further split Hymenoptera into bees (Anthophila, accounting for approximately 91% of observations) and non-bee hymenopterans (wasps and ants), on account of their differences in use of floral resources and foraging behaviour (Borchardt *et al*., 2024; Zemenick *et al*., 2019), resulting in five pollinator groups.

To characterise visitation patterns, we employed Principal Component Analysis (PCA) on the proportions of visits by each of the five pollinator groups to each plant species. This method provided orthogonal variables appropriate for testing associations while minimising collinearity (Mundry, 2014), and allowed us to leverage the continuous variation in visitation patterns, rather than artificially grouping plant species into discrete pollinator categories. The first three principal components (PC1–PC3) explained 84.5% of total variation, with PC1 (41.1%) clearly separating bee- vs. fly-dominated visitation (**Figure 3A and B**).

### Floral display trait categories

As part of the floral display, we considered flower shapes and colour. We classified all 43 species into five shape categories (bilabiate, funnel, head, tube, disk) following Pellissier *et al*. (2010) and Fantinato *et al*. (2016) (**Table S1**). Bilabiate and disk flowers represent the extremes in a gradient of floral morphological specialisation (higher to lower), while the other categories exhibit intermediate positions.

Floral colour categories were estimated using floral reflectance spectra and bee visual models. Flower reflectance was measured between 300 and 700 nm using a FLAME-T-UV-VIS Spectrometer and a Pulsed Xenon Lamp (Ocean Insight), using a probe holder at a 45° angle to the tissue surface. In total, 38 species were measured in this way (mean n = 4.5). In addition, spectral data for 5 species were obtained from the Floral Reflectance Database (Arnold *et al*., 2008), which was measured using a similar protocol (Chittka *et al*., 1994). A comparison of 22 species overlapping across these datasets showed comparable reflectance curves (**Figure S1A**). Colours in the bee visual space were employed as a more biologically meaningful categorisation than human vision. Bee-colours were reconstructed with the R package “pavo” (Maia *et al*., 2019), and the built-in *Apis mellifera* model. The specific parameters for the bee visual model are provided in **Methods S2**. The median distance in the bee colour space for species overlapping between datasets was 0.05 hexagon units (**Figure S1B**), where 0.11 units represent the commonly accepted limit of colour discrimination for non-trained bees in experimental studies (Dyer, 2006; Paine *et al*., 2019). Colour categories correspond to the sections of the bee-visual colourspace hexagon (bee-colour categories: “green”, “uv-blue”, “blue”, “uv-green” and “blue-green”; **Table S1**).

### Statistical analyses

All statistical analyses were carried out using R (R Core Team, 2021), version 4.4.2 (2024-10-31).

#### Phylogenetic tree and phylogenetic signal tests

We tested for phylogenetic signal in two aspects of nectar composition: in the occurrence of individual metabolites and in the mean values of aggregated nectar traits. Additionally, we considered phylogenetic structure as a potential driver of variation in statistical models examining the associations among nectar traits, visitation patterns, and floral display traits (details in the next section).

We retrieved a phylogenetic tree for plant species from the synthetic tree available in the Open Tree of Life database (OpenTreeOfLife *et al*., 2020) (version 14.9) using the package rotl (Michonneau *et al*., 2016). The resulting subtree contained no polytomies, and branch lengths were estimated following Grafen’s method (Grafen, 1989) as implemented by the package APE (Paradis & Schliep, 2019), using power = 1. Note that *Phyteuma orbiculare* appears inside the *Campanula* clade because the latter genus is reported to be polyphyletic (Xu & Hong, 2021).

To assess the phylogenetic signal of individual metabolites, given the high proportion of 0 values, we used the D statistic (Orme *et al*., 2023) on species-level presence/absence values (where the detection of a metabolite in at least one sample was considered as presence for the species). For aggregated nectar traits, we evaluated their phylogenetic signal (Orme *et al*., 2023) using Pagel’s λ on species-average trait values.

#### Association between nectar traits, insect visitation and floral display traits

To evaluate associations between nectar traits and either visitation patterns or floral display categories, we used Generalised Linear Mixed Models (GLMMs) via glmmTMB (Brooks *et al*., 2017). All models examined nectar trait variation at the sample level (individual plants; n = 285 overall; n = 248 for models that integrated visitation data), with plant species as a random factor. To account for potential phylogenetic effects, extended models were employed, in which a phylogenetic distance matrix (branch length) was used as a covariance structure using the “propto” function (Brooks *et al*., 2017). Model selection between simple and phylogenetically informed models was based on the Akaike Information Criterion (AIC). Model assumptions were verified by visual evaluation of simulated residuals via DHARMa (Hartig, 2024). The significance of multiple comparisons in models evaluating floral display categories was evaluated with the multcomp package (Hothorn *et al*., 2008). To visualise associations between nectar traits and visitation principal components (PCs), we estimated marginal mean predictor effects using the emmeans package (Lenth & Piaskowski, 2026).

To further explore the association between nectar metabolomic profiles, visitation patterns as captured by PC1, and plant phylogeny, we performed a distance-based redundancy analysis (dbRDA; Legendre & Anderson, 1999) via the vegan package (Oksanen *et al*., 2025). To identify the metabolites driving the observed relationship between profiles and visitation PC1, we further conducted vector fitting using the envfit function (Oksanen *et al*., 2025).

In the case of models evaluating the association between visitation PCs and floral display categories (shape and colour categories, tested separately), these were fitted using generalised least squares from nlme (Pinheiro *et al*., 2026; Pinheiro & Bates, 2000), considering plant species-level data (n = 35). Plant phylogeny was integrated in these models via a correlation structure obtained with corPagel (Paradis & Schliep, 2019), with a non-fixed λ transformation parameter. Multiple comparisons among categories were performed via Tukey contrasts (Hothorn *et al*., 2008).

To account for biases in nectar compound richness and compositional dissimilarity arising from instrumental detection limits (Petrén *et al*., 2024; Wetzel & Whitehead, 2020), we included per-sample nectar volume as a continuous covariate. To both mitigate detection-limit artifacts and stabilise model fitting under various distribution families, samples collected via nectar washing were assigned a constant volume offset smaller than the minimum measured volume in the dataset. Sensitivity testing confirmed that the choice of this offset value did not impact the statistical significance of our models. To further confirm that technical detection limits did not drive the observed associations for nectar richness and dispersion variables, we explored alternative models using the total amount of the main sugars (sucrose, fructose and glucose, from FID quantification) per sample as a covariate, instead of sample volume. These alternative models identified the same associations with very similar significance levels (Tables S12-14).

## Results

### Nectar presented highly diverse metabolomes

We screened 70 metabolic compounds across all 43 species using untargeted mass spectrometry (MS) analysis, with identification to at least the chemical class level (**Figure 1, coloured circles, and Table S3**). These metabolites were categorised into eight chemical classes: sugars, amino acids, fatty acids, phytohormones, other primary metabolites and compounds implicated in signalling interactions, defensive compounds, and others. Sugars and related compounds dominated this dataset, comprising 42 compounds and representing more than half of the identified metabolomic profile. We identified 4 amino acids (phenylalanine, 2-aminoadipic acid, glycine and proline), seven fatty acids (including hexanoic acid, octanoic acid, nonanoic acid, palmitic acid, stearic acid), and 9 other primary metabolites. Additionally, we putatively identified single representatives of a phytohormone (1-indole-3-acetic acid, IAA) and a potential bee-attractant (2-octene-2-ol) (Simon *et al*., 2021), alongside four defensive compounds (quininic acid, oxalic acid, phenol and 2-acetylcyclopentanone). Among these metabolites, 28 compounds showed significant phylogenetic signals (compound codes in bold in **Figure 1**, and detailed in **Table S3**).

Given the importance and high abundance of sugar-related compounds, we also quantified the most abundant ones using flame ionisation detection (FID), a complementary method better suited to abundant compounds. This resulted in 22 compounds (**Figure 1**, grey-shaded circles), largely overlapping with the ones detected by the MS approach. The three common nectar sugars (fructose, glucose and sucrose) were present in all the sampled species. The other 19 sugar-related compounds (i.e., uncommon nectar sugars) were found in subsets of the total species, and comprised 11 disaccharides, 2 monosaccharides and 6 sugar alcohols. We found no phylogenetic signal in the quantities of the three main sugars (Pagel’s λ associated *p* >0.05). In contrast, the presence/absence of uncommon nectar sugars showed significant phylogenetic signal in 11 of 19 cases (compound codes in bold in **Figure 1**; detailed in **Table S3**).

### Nectar aggregated traits showed no phylogenetic signal

Despite the phylogenetic signals found in several individual nectar components, we did not detect a phylogenetic signal in the aggregated nectar traits (sucrose proportion, metabolomic richness, uncommon sugar richness, metabolomic dispersion, or uncommon sugar dispersion; **Figure 2, Table S4**).

**Figure 2:**
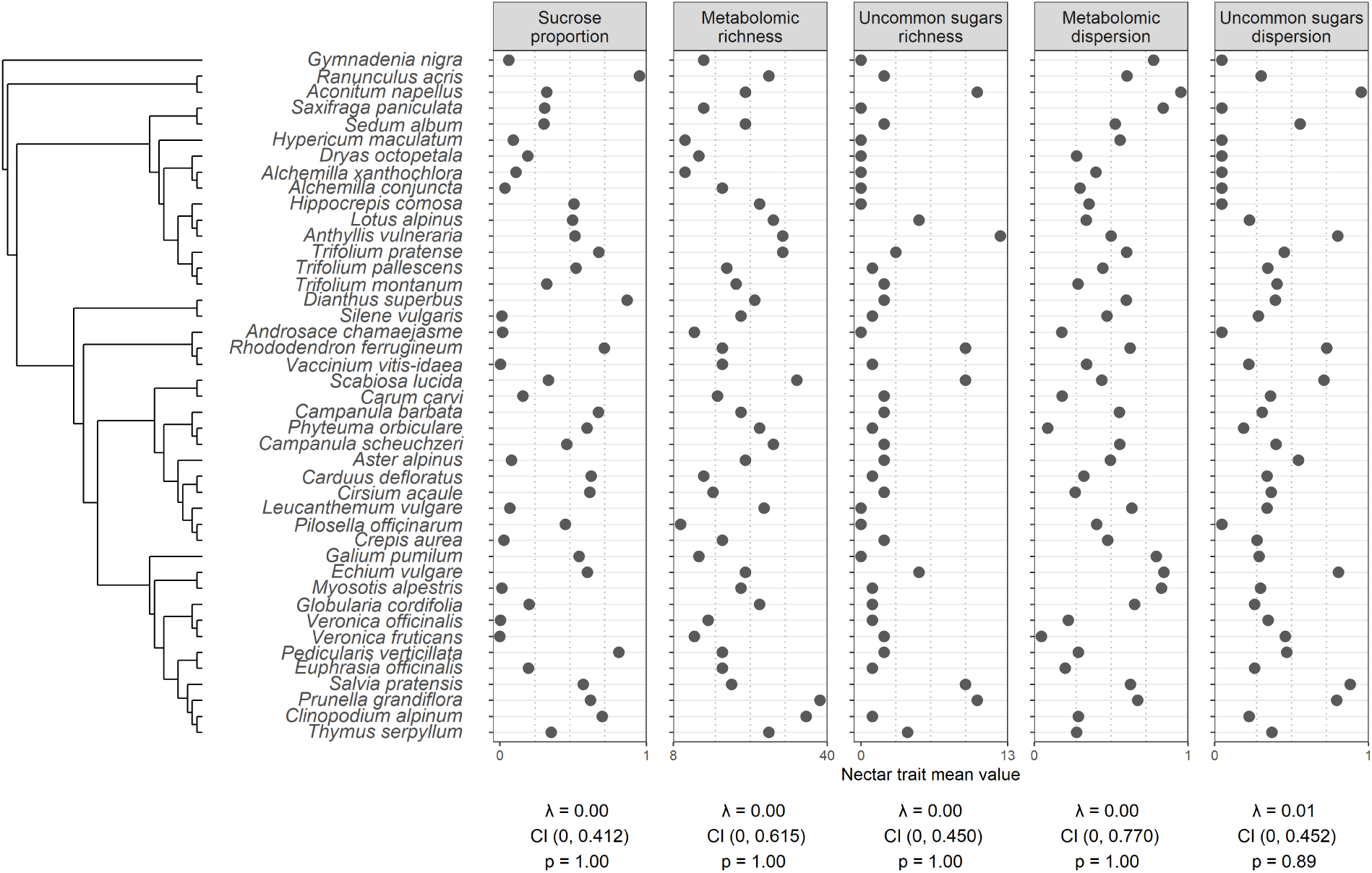
Nectar aggregated traits across the phylogenetic tree of species in our study system. Pagel’s λ values displayed at the bottom show the phylogenetic signal of each trait, together with its 95% Confidence Interval (CI) and associated p-value.

### Nectar traits correlated with insect visitation patterns

Across a total of 975 visitation events for 35 plant species, bees (Anthophila) and flies (Diptera) accounted for 35.5% and 33.6% of all visits, respectively. Lepidoptera visits contributed another 21.0% of total visits, nearly half of which were from *Adscita* spp. moths, while coleopterans represented 6.7% of all records. To characterise the overall visitation patterns, we employed a PCA on per-species visitation proportions across these pollinator groups (**Figure 3A**). This revealed that PC1 (41.1% of variation) clearly separated bee-dominated visitation (positive values) from fly-dominated visitation (negative values, **Figure 3B**). PC2 (24.3%) mostly separated lepidopteran-visited species, and PC3 (19.1%) contrasted visitation by most groups (except bees) from dipteran-visited species.

**Figure 3:**
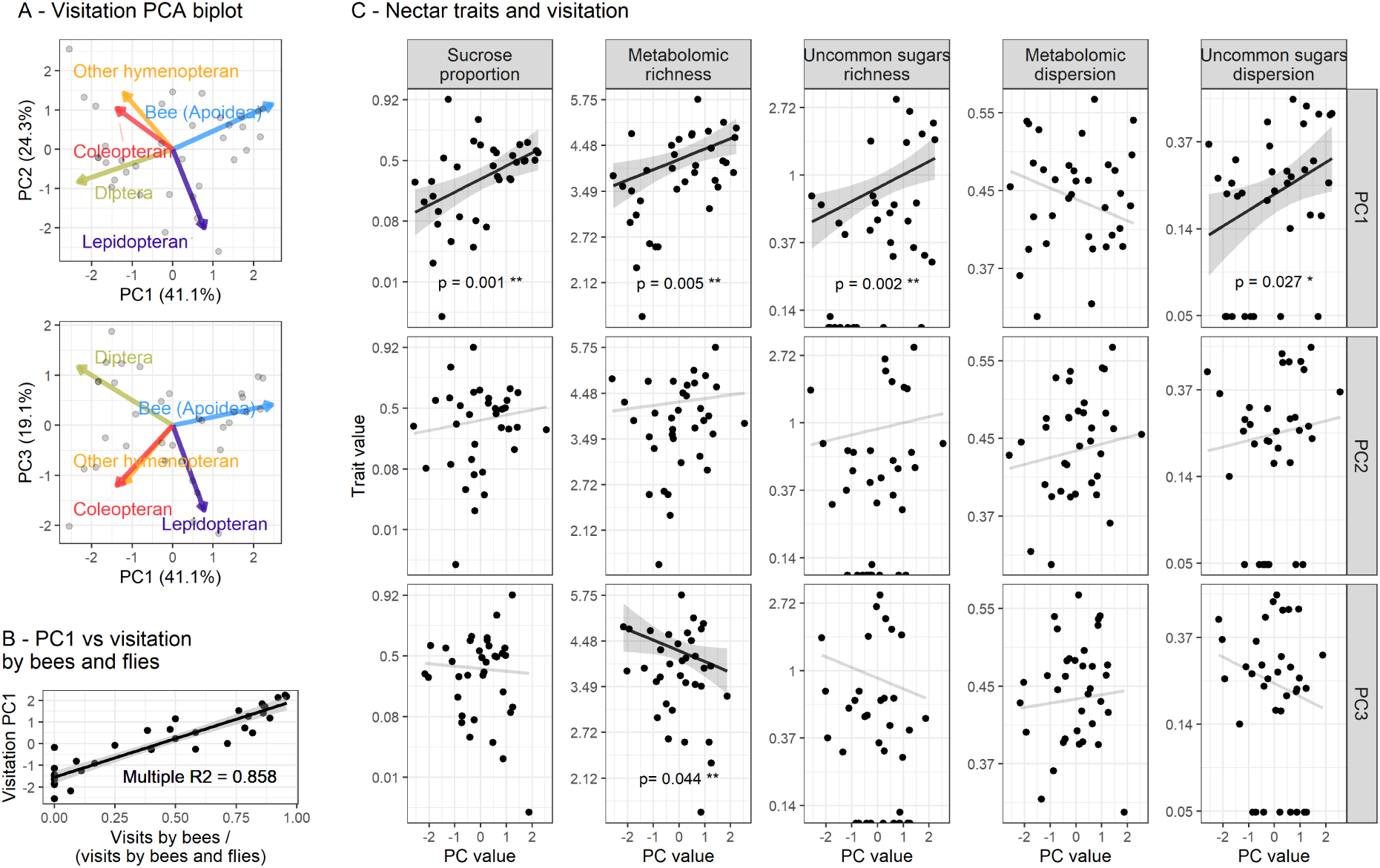
Patterns of insect visitation to plant species and their relationship to aggregated nectar traits. **A**. Biplot of the first three Principal Components (PCs) of the observed floral visitation, showing loading scores as arrows. **B**. Relationship between PC1 and the proportion of visitation by bees to that of bees plus flies, showing a high correlation in these metrics. **C**. Relationship between aggregated nectar traits and visitation PCs. For visualisation, points represent species-level means (n = 35), and trendlines and confidence intervals represent predictor estimates from visitation patterns from GLMMs (see methods section for details). Significant relationships (*p* < 0.05) are shown in black (full models in **Table S5**). The axes were transformed for visualisation, with sucrose proportion logit-transformed and other nectar traits log-transformed

Most aggregated nectar chemical traits were positively associated with the visitation PC1, reflecting an increase in these traits as visitation shifted from fly- to bee-dominated, including sucrose proportion (odds ratio = 1.71, *p* = 0.001), metabolomic richness (rate ratio = 1.06, *p* = 0.005), uncommon sugars richness (rate ratio = 1.22, *p* = 0.002), and multivariate dispersion in the uncommon sugars profile (rate ratio = 1.16, *p* = 0.027; **Figure 3C**). No significant associations were found between nectar traits and visitation PC2, whereas metabolomic richness was negatively associated with PC3 (rate ratio = 0.94, *p* = 0.044), indicating that species with higher visitation by lepidopterans/coleopterans/non-bee hymenopterans (as opposed to dipterans) had a slightly higher metabolomic richness. The site and sample volume were included as model covariates (the latter because richness and dispersion estimates can be inherently sensitive to sampling effort). Of these, the higher-elevation site had a negative effect on the proportion of sucrose and metabolomic dispersion (*p* <0.001 and = 0.008, respectively), while sample volume was positively associated with richness and dispersion variables (*p* <0.001). All full models can be found in **Table S5**.

In addition to the aggregated nectar trait tested above, we examined whether the full nectar metabolome profile (using presence/absence of metabolites) varied along visitation PC1. The phylogenetically informed dbRDA model (n = 35, *p* = 0.001, overall ecological model; adjusted *R*^2^ = 0.083) showed that phylogenetic structure (conditioned covariates) accounted for 16.47% of the total variation in metabolome profile, while the other two predictors (visitation PC1 and maximum nectar volume analysed per species) jointly explained an additional 8.40% of the variance. Marginal permutation tests revealed that visitation PC1 was a significant driver of metabolomic composition (*F*_1,27_ = 2.26, *p* = 0.002), whereas the maximum analysed nectar volume per species was not (*F*_1,27_ = 0.99, *p* = 0.459; models in **Table S6**).

We also identified which metabolites correlated most strongly with the constrained dbRDA ordination space (driven by visitation PC1 and maximum nectar volume) using vector fitting as a post hoc test. A total of 22 metabolites were significantly correlated with this ordination space (p <0.05, **Table S7**). Because visitation PC1 was the sole significant constraint in the model, these vectors capture compound alignment along the visitation gradient. The 10 most strongly correlated metabolites (p <0.002, **Figure 4A**) included 7 sugars (4 monosaccharides and 3 disaccharides), 2 fatty acids (putatively stearic acid and nonanoic acid) and one primary metabolite (putatively xylopyranose). These metabolites were distributed across the phylogenetic tree (**Figure 4B**), and their presence was consistently associated with *positive* values of PC1 (bee-dominated visitation). Conversely, there were no metabolites that significantly associated with negative PC1 values (fly-dominated visitation).

**Figure 4:**
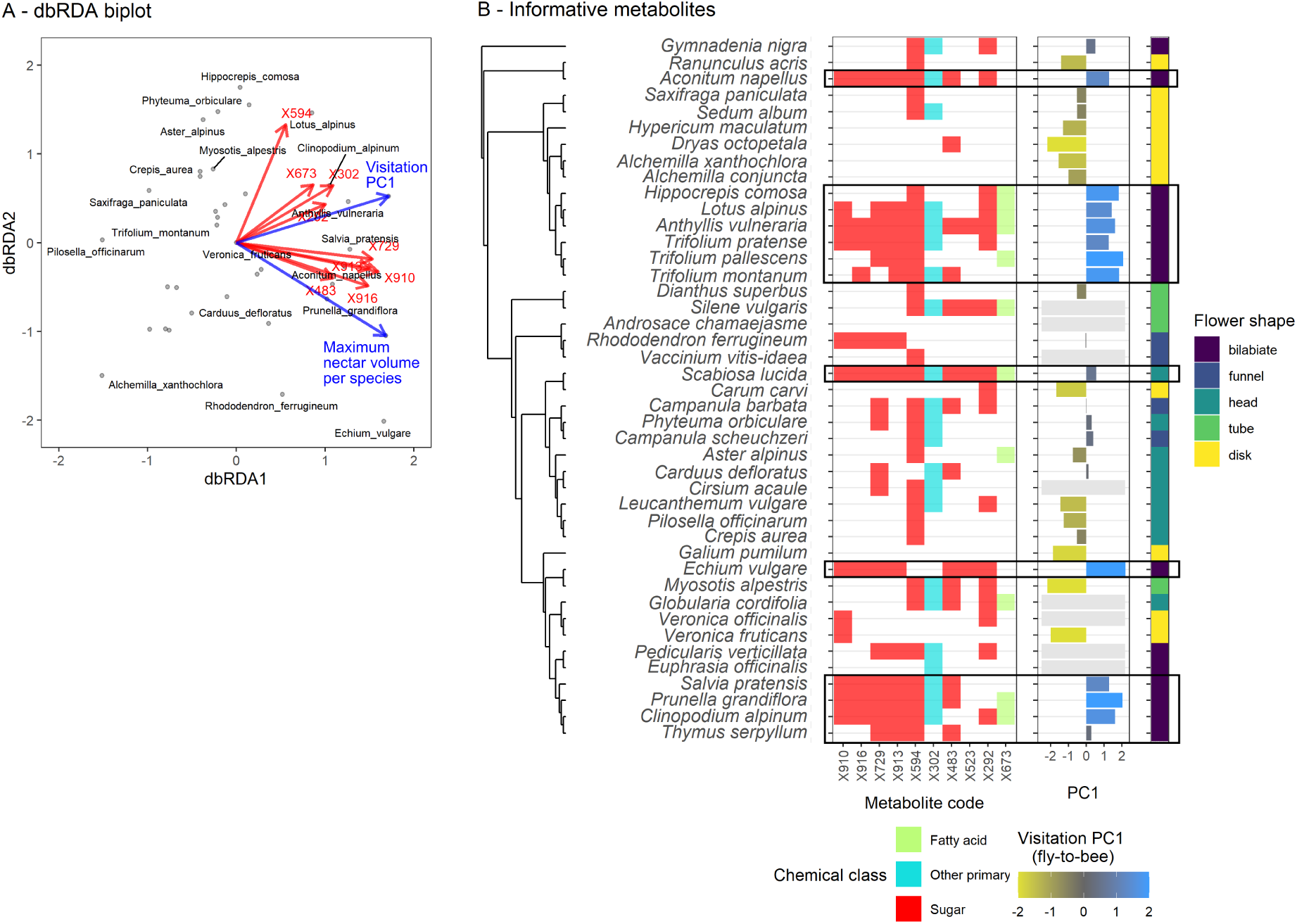
Association of insect visitation patterns with nectar metabolomes. **A**. Phylogeny- informed distance-based Redundancy Analysis (dbRDA) biplot, showing that the presence of individual metabolites (red arrows, p value <0.005) was related to visitation PC1 (blue arrow), which was mainly driven by bee vs fly visitation (positive vs negative values, respectively). **B**. Distribution of the 10 most informative metabolites for the dbRDA, as occurring across the 43 plant species (presence is shown as solid squares). The visitation PC1 (displayed for the 35 species with available data) and floral shape are depicted on the center-left and left, respectively. Note that species with a high proportion of metabolite presence (highlighted black boxes) often show high values of visitation PC1 and bilabiate flowers.

### Floral display traits reflected visitation patterns

Both flower shape and colour were associated with fly-bee visitation patterns (**Figure 5A**). The most complex flower shape (bilabiate) showed higher PC1 values than the simplest shape (disk), with other flower-shape categories falling in between (full models in **Table S8**). For colour, ultraviolet-blue and blue flowers showed significantly higher PC1 values than blue-green ones (full models in **Table S9**).

**Figure 5:**
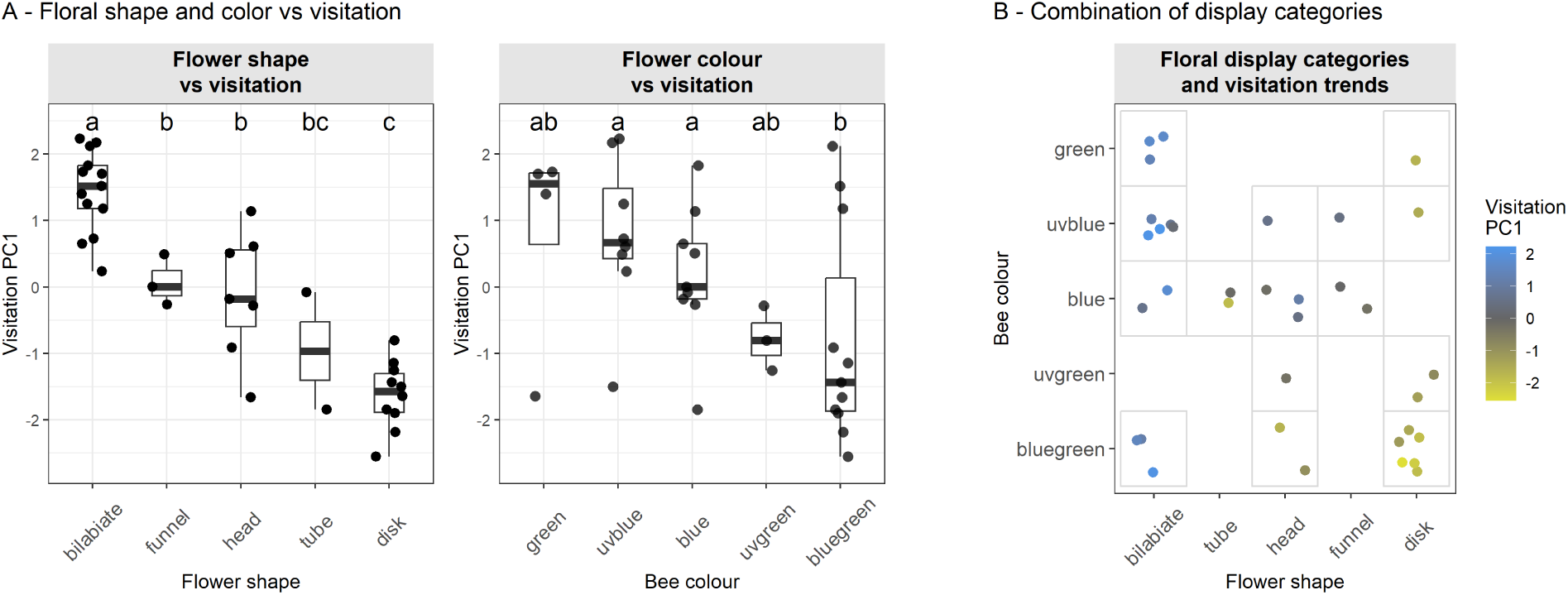
Flower display categories and their relationship with visitation. **A.** Visitation PC1 values of species grouped by their flower shape and colour category. The letters above bars indicate groups with significant differences (full models in **Table S8** and **S9**). **B.** The combinations of both of these categories across all species in our study system.

Regarding potentially coordinated display strategies, 7 species combined bilabiate shape with ultraviolet-blue or blue colour (both types of categories most preferred by bees), while 6 species combined disk shape with blue-green colour (both categories most preferred by flies). Thus, almost one third of the species (13/43) exhibited one of these two “polar opposite” combinations of traits. Several other species displayed mixed shape–colour combinations, with many of these falling roughly in the negative diagonal, although this pattern was not universal (**Figure 5B**).

### Nectar chemical richness traits were associated with floral display

Overall, nectar aggregated traits showed similar associations with flower shape and colour categories (**Figure 6**, top and lower row, respectively), although these associations were not consistently significant across traits. Specifically, nectar metabolomic richness and uncommon sugar richness were highest in bilabiate flowers and lowest in disk flowers and were also higher in blue or uv-blue flowers than in blue-green ones (full models in Tables S10 and S11). These flower categories were also the most (bilabiate, blue/uv-blue) and least (disk, blue-green) preferred by bees (previous section). The other nectar aggregated traits, including sucrose proportion and dispersion variables, showed no significant differences.

**Figure 6:**
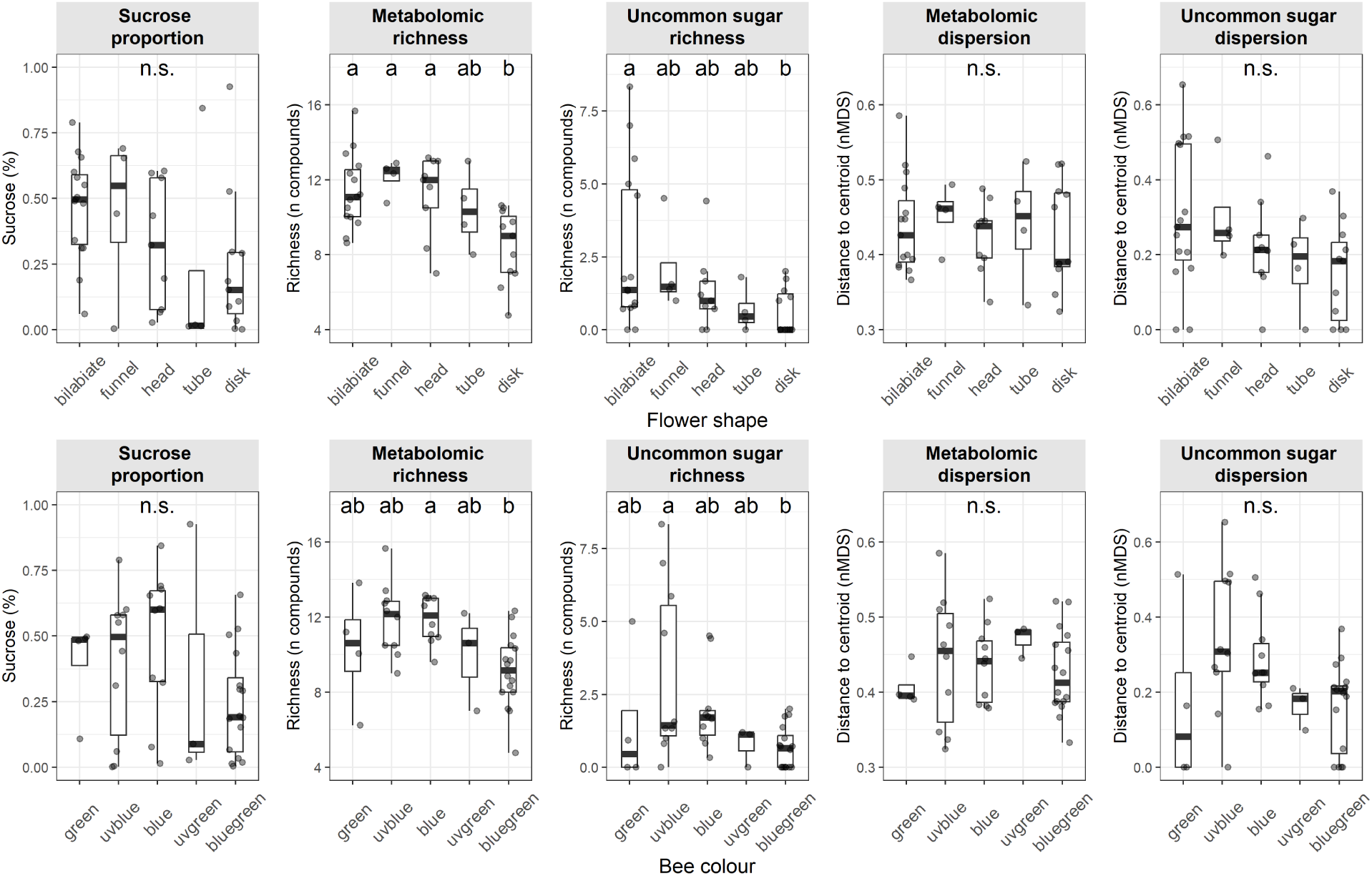
Association between aggregated nectar traits and floral display categories: shape (top row) and colour (bottom row). In each panel, the *y* represents species-average values for nectar traits, while floral display categories are shown in the *x* axis. The letters above bars indicate groups with significant differences (p < 0.05; full models in **Table S10** and **S11**).

## Discussion

Multiple aggregated nectar chemical traits tracked the most prominent insect visitation gradient in our system, including sucrose proportion, metabolomic richness, and uncommon sugar richness and dispersion. Higher values of each of these traits were associated with bee visitation and lower values with fly visitation independently of plant phylogeny. Underpinning the association between chemical richness and visitation, we identified multiple individual metabolites whose presence was consistently associated with bee visitation in separate taxa across the phylogenetic tree.

Conversely, no metabolites were distinctly associated with fly visitation. We also found that nectar chemical traits were sometimes linked to floral display traits such as flower colour and shape, suggesting partial phenotypic integration among these floral phenotypic components. Together, these findings provide evidence that complex features of nectar chemistry play an important functional role in mediating plant–pollinator interactions in natural communities.

### Individual compounds and aggregate nectar traits present different constraints

Nectar chemistry is a complex phenotype, encompassing both individual compounds and the compositional, aggregate-level properties they produce (such as chemical richness and dispersion). Similarly to the chemistry of plant vegetative tissues, these components can be subject to diverse selection pressures and show phylogenetic and metabolic constraints (Kessler & Kalske, 2018).

Consistent with previous studies (Nicolson & Thornburg, 2007; Palmer-Young *et al*., 2019), we found that individual compounds in nectar were highly variable among species. Many of these individual components also showed phylogenetic signals in their distribution, indicating phylogenetic or metabolic constraints. However, similar to other studies on multi-compound nectar traits in co-occurring species (Venjakob *et al*., 2022), we found no phylogenetic signal in any of the aggregated nectar traits. By contrast, cross-species comparisons from studies not focused on species that co-occur within natural communities have identified weak phylogenetic signals in nectar traits (Chalcoff *et al*., 2017). This suggests that these patterns are scale-dependent and that local ecological processes within communities contribute to shaping nectar trait evolution (Losos, 2008).

In contrast to the phylogenetic signal found in individual metabolites, we found that aggregate properties of nectar composition carry a phylogeny-independent signal of pollinator-associated convergence. A comparable pattern has been reported for floral scent: in *Campanula*, the overall scent bouquet showed no phylogenetic signal, whereas several individual volatile constituents did (Milet-Pinheiro *et al*., 2021). In principle, aggregated traits such as nectar metabolomic richness and dispersion can be reached through many different combinations of compounds, each with their own phylogenetic or metabolic constraints. This allows different lineages to converge on similar aggregated nectar traits — whether as a direct target of pollinator-mediated selection or through covariation with selected metabolites.

### Nectar chemistry suggests convergent pollination strategies

Multiple nectar chemical traits tracked the bee-fly visitation gradient across our alpine system independently of phylogeny, supporting nectar aggregated traits as convergent, pollinator-associated components of the floral phenotype. The main axis of variation in floral visitation distinguished bee- from fly-visited plant species, with these two insect groups being the most prevalent pollinators in alpine systems (Arroyo *et al*., 1982; Kearns, 1992; Lázaro *et al*., 2008; Lefebvre *et al*., 2018). Aggregated nectar traits were associated with this axis, with bee-visited plants showing a higher sucrose proportion, chemical richness (for both overall metabolome and uncommon sugars) and dispersion in uncommon sugar profiles. Patterns in sucrose proportion were consistent with previous findings (Abrahamczyk *et al*., 2017; Chalcoff *et al*., 2017; Liu *et al*., 2024), but, to our knowledge, our study represents the first empirical evidence that nectar phytochemical diversity is linked to visitation by different pollinator functional groups.

This result matches broader patterns in which plant chemical richness mediates biotic interactions, including the deterrence of non-specialised consumers (Berenbaum & Zangerl, 1996; Kessler & Kalske, 2018). Indeed, the metabolites associating most strongly towards bee visitation recurred across multiple, distantly related clades, consistent with convergent evolution. However, the high turnover of these metabolites across species (which show a mosaic pattern of presence/absence) points toward some redundancy in their individual contribution to this function, supporting the hypothesis that compound richness, rather than the identity of any single compound, may be regarded as the more functionally relevant axis of nectar complexity in our system. A recent study by MacNeill *et al*. (2025) on plants specialised in two nectar consumers (bees and hummingbirds) also found that certain metabolite groups in nectar were associated with these pollination types. In our system, flies provided a different type of contrast, as they are comparatively opportunistic and generalised visitors (Abrahamczyk *et al*., 2017; Oldroyd, 1964; Tiusanen *et al*., 2016), and thus their preferences may be limited by multiple co-occurring metabolites additively or synergistically lowering nectar palatability (Berenbaum & Zangerl, 1996). These patterns notwithstanding, our observational design cannot distinguish whether these compounds actively attract bees, deter flies, or both: because pollinator visitation was characterised as a relative (compositional) gradient, an increase in bee-associated chemistry is mathematically inseparable from a decrease in fly-associated visitation. Future experiments with choice-based bioassays would be necessary to disentangle these non-mutually-exclusive mechanisms.

### Nectar chemistry and floral display traits suggest partial phenotypic integration

Despite the clear association between nectar chemistry and pollinator visitation, the associations between nectar traits and floral display traits (shape and colour) were only significant in the case of compound richness. The pollination syndrome hypothesis predicts coordinated evolution of multiple floral traits in response to shared pollinator selection (Faegri & Van Der Pijl, 1979; Rosas-Guerrero *et al*., 2014). Our results suggest that, at least in this system, nectar chemistry does not evolve as tightly coupled to visual display as might be expected under tight phenotypic integration. A similar pattern has been reported in Polemoniaceae, where Rose & Sytsma (2021) found that floral trait evolution converged with pollinator type while being only partially explained by co-variation within floral traits. Theoretical work further suggests that moderate modularity among floral trait suites, rather than tight integration, can facilitate adaptive flexibility under shifting pollinator selection (Dellinger *et al*., 2019). Moreover, strong integration is most expected under self-pollination or high insect specialisation (Rosas-Guerrero *et al*., 2011). Neither of these modes characterise alpine communities, typically highly reliant on insect pollination (Arroyo *et al*., 1982; Körner, 1999) with a generalist-specialist pollination spectrum by multiple insect guilds.

These findings also raise a question about the biogeographical distributions of floral traits: in alpine environments, bee and fly abundance are known to shift consistently with elevation (Kearns, 1992; Lefebvre *et al*., 2018; Sommaggio *et al*., 2022). In the Alps, flies become the dominant pollinator group above 1500 m (Lefebvre *et al*., 2018; Sommaggio *et al*., 2022).

Simultaneously, simpler floral morphologies also tend to predominate at high elevations (Fantinato *et al*., 2016; Pellissier *et al*., 2010). Whether nectar phytochemical diversity declines similarly with elevation, mirroring the chemical and morphological pattern we report here for the bee-fly visitation gradient, remains an open but tractable question, potentially relevant to the altitudinal niche breadth hypothesis (Rasmann *et al*., 2014).

### Closing remarks

While previous work has established that individual metabolites in nectar can play a functional role in shaping plant-pollinator interactions (Barberis *et al*., 2023; Gegear *et al*., 2007; Hansen *et al*., 2007; Nepi, 2014; Nicolson & Thornburg, 2007; Wong *et al*., 2023; Wright *et al*., 2013), our findings strongly suggest that aggregate-level features of nectar composition — including phytochemical diversity — are also ecologically important. By evaluating nectar metabolomic richness and dispersion, we provide evidence for convergent chemical strategies across disparate taxa visited by the same pollinator groups. While more work is needed to extend these observations to other systems, our results suggest broad patterns that may hold generally, such as a tendency for relatively simple nectar profiles — perhaps in association with simple floral shapes — to mediate visitation by broadly generalist pollinator groups, such as flies, while more complex chemical traits and floral structures reflect visitation by specialist pollinators, including bees.

Integrating the chemistry of floral rewards into the study of floral phenotypes and interactions in nature also provides an avenue to understand novel aspects of pollination and community ecology. For instance, the association between floral traits and visitation described above echoes the concept of a generalised pollination syndrome, which has historically found its characterisation elusive (Dellinger, 2020; Ollerton *et al*., 2007). In this regard, our results suggest that generalised pollination syndromes may rely on chemical simplicity rather than specific chemical signatures.

Comparing floral reward traits within plant communities and identifying signatures of trait divergence and convergence may also help to explain plant community dynamics, such as reproductive facilitation, competition, and ultimately patterns of plant species coexistence (E-Vojtkó *et al*., 2020; Sargent & Ackerly, 2008).

## Acknowledgements

We thank Jake Alexander and Camille Brioschi for assistance with various logistical aspects of fieldwork in Calanda, and Florian Schiestl for sharing sugar standards. We also thank student assistants for their contribution in fieldwork, including Fabio Schwarzenbach, Dominic Stadler, Tomoki Loeillot, Ziyu Zhu and Paloma Juliá-Martínez. We thank Aphrodite Kantsa for helpful discussions on the study. RR, YY and PZ acknowledge the SNSF Prima Grant (PR00P3_193237). RR and PZ also acknowledge the RESPONSE Fund from the European Union’s Horizon 2020 research and innovation program under the Marie Skłodowska-Curie(No. 547585). YY was supported by the Hainan Provincial Natural Science Foundation of China (323QN196) and the China Scholarship Council (202107565020).

## Statement of authorship

RR and PZ conceived and designed the study. RR, LS and YY conducted fieldwork. RR, LS, YY and JS conducted laboratory work and chemical analyses. RR analysed the data. RR and PZ drafted the manuscript. All authors contributed critically to manuscript revisions. No conflicts of interest exist.

## Data accessibility statement

The data that support the findings of this study will be made openly available upon publication.

## Contents of supporting information

- Table S1: List of plant species sampled
- Methods S1: Extraction and metabolomic analysis
- Table S2: Summary table of the observed floral visitation
- Methods S2: Parameters in the bee colour visual model
- Figure S1: Validation of colour data from both data sources
- Table S3: List of metabolic compounds found
- Table S4: Phylogenetic signal on aggregated nectar traits
- Table S5: Summaries of GLMMs on the relationship between nectar traits and visitation PC1
- Table S6: dbRDA of metabolomic profiles against visitation and phylogeny
- Table S7: dbRDA vector fitting
- Table S8: Summary of GLMMs on the relationship between the visitation PC1 and floral shape categories
- Table S9: Summary of GLMMs on the relationship between the visitation PC1 and floral colour categories
- Table S10: Summaries of GLMMs on the relationship between nectar traits and floral shape categories
- Table S11: Summaries of GLMMs on the relationship between nectar traits and floral colour categories
- Tables S12-14: Summaries of GLMMs corresponding to Tables S5, S10 and S11, but using total content of main sugars as a covariate, instead of sample volume.

## References

Abdala-Roberts, L. & Moreira, X. (2024). Effects of phytochemical diversity on multitrophic interactions. Current opinion in insect science, 64, 101228.

Abrahamczyk, S., Kessler, M., Hanley, D., Karger, D.N., Müller, M.P., Knauer, A.C. et al. (2017). Pollinator adaptation and the evolution of floral nectar sugar composition. Journal of Evolutionary Biology, 30, 112–127.

Alexander, J.M., Diez, J.M. & Levine, J.M. (2015). Novel competitors shape species’ responses to climate change. Nature, 525, 515–518.

Allsopp, M., Nicolson, S. & Jackson, S. (1998). Xylose as a nectar sugar: the response of cape honeybees, *Apis mellifera capensis* eschscholtz (Hymenoptera: Apidae). African Entomology, 6, 317–323.

Apland, J.S. & Koski, M.H. (2025). Isolating the effects of floral temperature on visitation and behaviour of wild bee and fly pollinators. Functional Ecology, 39, 2496–2508.

Arnold, S., Savolainen, V. & Chittka, L. (2008). Fred: the floral reflectance spectra database. Nature Precedings, pp. 1–1.

Arnold, S.E., Savolainen, V. & Chittka, L. (2009). Flower colours along an alpine altitude gradient, seen through the eyes of fly and bee pollinators. Arthropod-Plant Interactions, 3, 27–43.

Arroyo, M.T.K., Primack, R. & Armesto, J. (1982). Community studies in pollination ecology in the high temperate Andes of central Chile. i. pollination mechanisms and altitudinal variation. American journal of botany, 69, 82–97.

Baker, H.G. & Baker, I. (1983). Floral nectar sugar constituents in relation to pollinator type. In: Handbook of experimental pollination biology. Van Nostrand Reinhold Company, New York, NY, USA, p. 117–141.

Ballarin, C.S., Fontúrbel, F.E., Rech, A.R., Oliveira, P.E., Goés, G.A., Polizello, D.S. et al. (2024). How many animal-pollinated angiosperms are nectar-producing? New Phytologist, 243, 2008–2020.

Barberis, M., Calabrese, D., Galloni, M. & Nepi, M. (2023). Secondary metabolites in nectar-mediated plant-pollinator relationships. Plants, 12, 550.

Barker, R.J. (1977). Some carbohydrates found in pollen and pollen substitutes are toxic to honey bees. The Journal of nutrition, 107, 1859–1862.

Barker, R.J. & Lehner, Y. (1974). Acceptance and sustenance value of naturally occurring sugars fed to newly emerged adult workers of honey bees (*Apis mellifera* l.). Journal of Experimental Zoology, 187, 277–285.

Berenbaum, M.R. & Zangerl, A.R. (1996). Phytochemical diversity: adaptation or random variation? In: Phytochemical diversity and redundancy in ecological interactions (eds. Romeo, J.T., Saunders, J.A. & Barbosa, P.). Springer, New York, NY, USA, pp. 1–24.

Borchardt, K.E., Holthaus, D., Soto Méndez, P.A. & Toth, A.L. (2024). Debunking wasp pollination: Wasps are comparable to bees in terms of plant interactions, body pollen and single-visit pollen deposition. Ecological Entomology, 49, 569–584.

Brooks, M.E., Kristensen, K., van Benthem, K.J., Magnusson, A., Berg, C.W., Nielsen, A. et al. (2017). glmmTMB balances speed and flexibility among packages for zero-inflated generalized linear mixed modeling. The R Journal, 9, 378–400.

Chalcoff, V.R., Gleiser, G., Ezcurra, C. & Aizen, M.A. (2017). Pollinator type and secondarily climate are related to nectar sugar composition across the angiosperms. Evolutionary ecology, 31, 585–602.

Chittka, L., Shmida, A., Troje, N. & Menzel, R. (1994). Ultraviolet as a component of flower reflections, and the colour perception of hymenoptera. Vision research, 34, 1489–1508.

Dellinger, A.S. (2020). Pollination syndromes in the 21st century: where do we stand and where may we go? New Phytologist, 228, 1193–1213.

Dellinger, A.S., Artuso, S., Pamperl, S., Michelangeli, F.A., Penneys, D.S., Fernández-Fernández, D.M. et al. (2019). Modularity increases rate of floral evolution and adaptive success for functionally specialized pollination systems. Communications biology, 2, 453.

Dyer, A.G. (2006). Discrimination of flower colours in natural settings by the bumblebee species *Bombus terrestris* (hymenoptera: Apidae). Entomologia generalis, 28, 257.

E-Vojtkó, A., de Bello, F., Durka, W., Kühn, I. & Götzenberger, L. (2020). The neglected importance of floral traits in trait-based plant community assembly. Journal of Vegetation Science, 31, 529–539.

Faegri, K. & Van Der Pijl, L. (1979). Principles of pollination ecology. Pergamon press, Oxford, UK.

Fantinato, E., Giovanetti, M., Del Vecchio, S., Buffa, G. et al. (2016). Altitudinal patterns of floral morphologies in dry calcareous grasslands. Plant Sociology, 53, 83–90.

von Frisch, K. (1971). Bees: Their Vision, Chemical Senses, and Language. Rev - revised, 1 edn. Cornell University Press, Ithaca, NY, USA.

Gegear, R.J., Manson, J.S. & Thomson, J.D. (2007). Ecological context influences pollinator deterrence by alkaloids in floral nectar. Ecology letters, 10, 375–382.

Grafen, A. (1989). The phylogenetic regression. *Philosophical Transactions of the Royal Society of London. B*, Biological Sciences, 326, 119–157.

Hansen, D.M., Olesen, J.M., Mione, T., Johnson, S.D. & Müller, C.B. (2007). Coloured nectar: distribution, ecology, and evolution of an enigmatic floral trait. Biological Reviews, 82, 83–111.

Hartig, F. (2024). DHARMa: Residual Diagnostics for Hierarchical (Multi-Level / Mixed) Regression Models. R package version 0.4.7.

Hauri, K.C., Glassmire, A.E. & Wetzel, W.C. (2021). Chemical diversity rather than cultivar diversity predicts natural enemy control of herbivore pests. Ecological Applications, 31, e02289.

Herrera, C.M. (1987). Components of pollinator” quality”: comparative analysis of a diverse insect assemblage. Oikos, pp. 79–90.

Herrera, C.M. (1990). Daily patterns of pollinator activity, differential pollinating effectiveness, and floral resource availability, in a summer-flowering Mediterranean shrub. Oikos, pp. 277–288.

Heuckeroth, S., Damiani, T., Smirnov, A., Mokshyna, O., Brungs, C., Korf, A. et al. (2024). Reproducible mass spectrometry data processing and compound annotation in mzmine 3. Nature protocols, 19, 2597–2641.

Hothorn, T., Bretz, F. & Westfall, P. (2008). Simultaneous inference in general parametric models. Biometrical Journal: Journal of Mathematical Methods in Biosciences, 50, 346–363.

Kantsa, A., Raguso, R.A., Dyer, A.G., Olesen, J.M., Tscheulin, T. & Petanidou, T. (2018). Disentangling the role of floral sensory stimuli in pollination networks. Nature communications, 9, 1041.

Kearns, C.A. (1992). Anthophilous fly distribution across an elevation gradient. American Midland Naturalist, pp. 172–182.

Kessler, A. & Kalske, A. (2018). Plant secondary metabolite diversity and species interactions. Annual Review of Ecology, Evolution, and Systematics, 49, 115–138.

Körner, C. (1999). *Alpine plant life: functional plant ecology of high mountain ecosystems*. Springer, Berlin, Germany.

Larson, B., Kevan, P. & Inouye, D.W. (2001). Flies and flowers: taxonomic diversity of anthophiles and pollinators. The Canadian Entomologist, 133, 439–465.

Lázaro, A., Hegland, S.J. & Totland, Ø. (2008). The relationships between floral traits and specificity of pollination systems in three Scandinavian plant communities. Oecologia, 157, 249–257.

Lefebvre, V., Fontaine, C., Villemant, C. & Daugeron, C. (2014). Are empidine dance flies major flower visitors in alpine environments? a case study in the Alps, France. Biology letters, 10.

Lefebvre, V., Villemant, C., Fontaine, C. & Daugeron, C. (2018). Altitudinal, temporal and trophic partitioning of flower-visitors in alpine communities. Scientific reports, 8, 4706.

Legendre, P. & Anderson, M.J. (1999). Distance-based redundancy analysis: testing multispecies responses in multifactorial ecological experiments. Ecological monographs, 69, 1–24.

Lenth, R.V. & Piaskowski, J. (2026). emmeans: Estimated Marginal Means, aka Least-Squares Means. R package version 2.0.3.

Leonhardt, S.D. & Blüthgen, N. (2012). The same, but different: pollen foraging in honeybee and bumblebee colonies. Apidologie, 43, 449–464.

Leonhardt, S.D., Chui, S.X. & Kuba, K. (2024). The role of non-volatile chemicals of floral rewards in plant-pollinator interactions. Basic and Applied Ecology, 75, 31–43.

Lisec, J., Schauer, N., Kopka, J., Willmitzer, L. & Fernie, A.R. (2006). Gas chromatography mass spectrometry–based metabolite profiling in plants. Nature protocols, 1, 387–396.

Liu, Y., Dunker, S., Durka, W., Dominik, C., Heuschele, J.M., Honchar, H. et al. (2024). Eco-evolutionary processes shaping floral nectar sugar composition. Scientific Reports, 14, 13856.

Losos, J.B. (2008). Phylogenetic niche conservatism, phylogenetic signal and the relationship between phylogenetic relatedness and ecological similarity among species. Ecology letters, 11, 995–1003.

MacNeill, F.T., Hunter, S.G., Muth, F. & Sedio, B.E. (2025). Nectar metabolomes contribute to pollination syndromes. New Phytologist, 247, 951–967.

Maia, R., Gruson, H., Endler, J.A. & White, T.E. (2019). pavo 2: new tools for the spectral and spatial analysis of colour in r. Methods in Ecology and Evolution, 10.

Michonneau, F., Brown, J.W. & Winter, D.J. (2016). rotl: an r package to interact with the open tree of life data. Methods in Ecology and Evolution, 7, 1476–1481.

Milet-Pinheiro, P., Santos, P.S.C., Prieto-Benítez, S., Ayasse, M. & Dötterl, S. (2021). Differential evolutionary history in visual and olfactory floral cues of the bee-pollinated genus *Campanula* (campanulaceae). Plants, 10, 1356.

Mundry, R. (2014). Statistical issues and assumptions of phylogenetic generalized least squares. In: Modern phylogenetic comparative methods and their application in evolutionary biology: Concepts and practice (ed. Garamszegi, L.Z.). Springer, pp. 131–153.

Nepi, M. (2014). Beyond nectar sweetness: the hidden ecological role of non-protein amino acids in nectar. Journal of Ecology, 102, 108–115.

Nicolson, S.W. (2022). Sweet solutions: nectar chemistry and quality. Philosophical Transactions of the Royal Society B, 377, 20210163.

Nicolson, S.W. & Thornburg, R.W. (2007). Nectar chemistry. In: Nectaries and necta*r* (eds. Nicolson, S.W., Nepi, M. & Pacini, E.). Springer, pp. 215–264.

Oksanen, J., Simpson, G.L., Blanchet, F.G., Kindt, R., Legendre, P., Minchin, P.R. et al. (2025). vegan: Community Ecology Package. R package version 2.7-0, https://github.com/vegandevs/vegan.

Oldroyd, H. (1964). The natural history of flies. Weidenfeld and Nicolson, London, UK.

Ollerton, J., Killick, A., Lamborn, E., Watts, S. & Whiston, M. (2007). Multiple meanings and modes: on the many ways to be a generalist flower. Taxon, 56, 717–728.

OpenTreeOfLife, Redelings, B., Reyes, L.L.S., Cranston, K.A., Allman, J., Holder, M.T., et al. (2020). Open tree of life synthetic tree.

Orme, D., Freckleton, R., Thomas, G., Petzoldt, T., Fritz, S., Isaac, N. et al. (2023). caper: Comparative Analyses of Phylogenetics and Evolution in R. R package version 1.0.3.

Paine, K.C., White, T.E. & Whitney, K.D. (2019). Intraspecific floral color variation as perceived by pollinators and non-pollinators: evidence for pollinator-imposed constraints? Evolutionary Ecology, 33, 461–479.

Palmer-Young, E.C., Farrell, I.W., Adler, L.S., Milano, N.J., Egan, P.A., Irwin, R.E. et al. (2019). Secondary metabolites from nectar and pollen: a resource for ecological and evolutionary studies. Ecology, 100.

Paradis, E. & Schliep, K. (2019). ape 5.0: an environment for modern phylogenetics and evolutionary analyses in R. Bioinformatics, 35, 526–528.

Pellissier, L., Pottier, J., Vittoz, P., Dubuis, A. & Guisan, A. (2010). Spatial pattern of floral morphology: possible insight into the effects of pollinators on plant distributions. Oikos, 119, 1805–1813.

Perret, M., Chautems, A., Spichiger, R., Peixoto, M. & Savolainen, V. (2001). Nectar sugar composition in relation to pollination syndromes in sinningieae (gesneriaceae). Annals of botany, 87, 267–273.

Petrén, H., Anaia, R.A., Aragam, K.S., Bräutigam, A., Eckert, S., Heinen, R. et al. (2024). Understanding the chemodiversity of plants: Quantification, variation and ecological function. Ecological Monographs, 94, e1635.

Pinheiro, J., Bates, D. & R Core Team (2026). nlme: Linear and Nonlinear Mixed Effects Models. R package version 3.1–169.

Pinheiro, J.C. & Bates, D.M. (2000). Mixed-Effects Models in S and S-PLUS. Springer, New York.

Polidori, C., Ferrari, A. & Ronchetti, F. (2025). Biology and behaviour of european wild bees. In: Hidden and Wild: An Integrated Study of European Wild Bees (eds. Cilia, G., Ranalli, R., Zavatta, L. & Flaminio, S.). Springer, pp. 49–118.

Power, E.F., Stabler, D., Borland, A.M., Barnes, J. & Wright, G.A. (2018). Analysis of nectar from low-volume flowers: A comparison of collection methods for free amino acids. Methods in ecology and evolution, 9, 734–743.

R Core Team (2021). R: A Language and Environment for Statistical Computing. R Foundation for Statistical Computing, Vienna, Austria.

Rasmann, S., Alvarez, N. & Pellissier, L. (2014). The altitudinal niche-breadth hypothesis in insect-plant interactions. Annual Plant Reviews: Insect-Plant Interactions, 47, 339–359.

Rosas-Guerrero, V., Aguilar, R., Martén-Rodríguez, S., Ashworth, L., Lopezaraiza-Mikel, M., Bastida, J.M. et al. (2014). A quantitative review of pollination syndromes: do floral traits predict effective pollinators? Ecology letters, 17, 388–400.

Rosas-Guerrero, V., Quesada, M., Armbruster, W.S., Pérez-Barrales, R. & Smith, S.D. (2011). Influence of pollination specialization and breeding system on floral integration and phenotypic variation in Ipomoea. Evolution, 65, 350–364.

Rose, J.P. & Sytsma, K.J. (2021). Complex interactions underlie the correlated evolution of floral traits and their association with pollinators in a clade with diverse pollination systems. Evolution, 75, 1431–1449.

Roy, R., Schmitt, A.J., Thomas, J.B. & Carter, C.J. (2017). Nectar biology: from molecules to ecosystems. Plant Science, 262, 148–164.

Salazar, D., Jaramillo, A. & Marquis, R.J. (2016). The impact of plant chemical diversity on plant–herbivore interactions at the community level. Oecologia, 181, 1199–1208.

Salazar, D., Lokvam, J., Mesones, I., Vásquez Pilco, M., Ayarza Zuñiga, J.M., de Valpine, P. et al. (2018). Origin and maintenance of chemical diversity in a species-rich tropical tree lineage. Nature Ecology & Evolution, 2, 983–990.

Sargent, R.D. & Ackerly, D.D. (2008). Plant–pollinator interactions and the assembly of plant communities. Trends in Ecology & Evolution, 23, 123–130.

Simon, S.J., Keefover-Ring, K., Park, Y.L., Wimp, G., Grady, J. & DiFazio, S.P. (2021). Characterization of Salix nigra floral insect community and activity of three native Andrena bees. Ecology and Evolution, 11, 4688–4700.

Sols, A., Cadenas, E. & Alvarado, F. (1960). Enzymatic basis of mannose toxicity in honey bees. Science, 131, 297–298.

Sommaggio, D., Zanotelli, L., Vettorazzo, E., Burgio, G. & Fontana, P. (2022). Different distribution patterns of hoverflies (diptera: Syrphidae) and bees (hymenoptera: Anthophila) along altitudinal gradients in Dolomiti Bellunesi National Park (Italy). Insects, 13, 293.

Stevenson, P.C. (2020). For antagonists and mutualists: the paradox of insect toxic secondary metabolites in nectar and pollen. Phytochemistry Reviews, 19, 603–614.

Stevenson, P.C., Nicolson, S.W. & Wright, G.A. (2017). Plant secondary metabolites in nectar: impacts on pollinators and ecological functions. Functional Ecology, 31, 65–75.

Tiedge, K. & Lohaus, G. (2017). Nectar sugars and amino acids in day-and night-flowering Nicotiana species are more strongly shaped by pollinators’ preferences than organic acids and inorganic ions. PLoS One, 12, e0176865.

Tiusanen, M., Hebert, P.D., Schmidt, N.M. & Roslin, T. (2016). One fly to rule them all—muscid flies are the key pollinators in the Arctic. Proceedings of the Royal Society B: Biological Sciences, 283.

Vandelook, F., Janssens, S., Gijbels, P., Fischer, E., Van den Ende, W., Honnay, O., et al. (2019). Nectar traits differ between pollination syndromes in balsaminaceae. Annals of Botany, 124, 269–279.

Väre, H., Lampinen, R., Humphries, C. & Williams, P. (2003). Taxonomic diversity of vascular plants in the European alpine areas. In: Alpine biodiversity in Europe. Springer, Berlin, Germany, pp. 133–148.

Venjakob, C., Ruedenauer, F., Klein, A.M. & Leonhardt, S. (2022). Variation in nectar quality across 34 grassland plant species. Plant Biology, 24, 134–144.

Wetzel, W.C. & Whitehead, S.R. (2020). The many dimensions of phytochemical diversity: linking theory to practice. Ecology letters, 23, 16–32.

Wong, D.C., Pichersky, E. & Peakall, R. (2023). Many different flowers make a bouquet: Lessons from specialized metabolite diversity in plant–pollinator interactions. Current opinion in plant biology, 73, 102332.

Wood, T.J., Ghisbain, G., Rasmont, P., Kleijn, D., Raemakers, I., Praz, C. et al. (2021). Global patterns in bumble bee pollen collection show phylogenetic conservation of diet. Journal of Animal Ecology, 90, 2421–2430.

Wright, G., Baker, D., Palmer, M., Stabler, D., Mustard, J., Power, E. et al. (2013). Caffeine in floral nectar enhances a pollinator’s memory of reward. science, 339, 1202–1204.

Xu, C. & Hong, D.Y. (2021). Phylogenetic analyses confirm polyphyly of the genus *Campanula* (campanulaceae s. str.), leading to a proposal for generic reappraisal. Journal of Systematics and Evolution, 59, 475–489.

Zemenick, A.T., Kula, R.R., Russo, L. & Tooker, J. (2019). A network approach reveals parasitoid wasps to be generalized nectar foragers. Arthropod-Plant Interactions, 13, 239–251.

