## Supplementary material for "Nectar chemistry reflects pollination strategies in alpine plant communities"

**Contents**

**Table S1**: List of plant species sampled

**Methods S1**: Extraction and metabolomic analysis

**Table S2**: Summary table of the observed floral visitation

**Methods S2**: Parameters in the bee colour visual model

**Figure S1**: Validation of colour data from both data sources

**Table S3**: List of metabolic compounds found

**Table S4**: Phylogenetic signal on aggregated nectar traits

**Table S5**: Summaries of GLMMs on the relationship between nectar traits and visitation PC1

**Table S6**: dbRDA of metabolomic profiles against visitation and phylogeny

**Table S7**: dbRDA vector fitting

**Table S8**: Summary of GLMMs on the relationship between the visitation PC1 and floral shape categories

**Table S9**: Summary of GLMMs on the relationship between the visitation PC1 and floral colour categories

**Table S10**: Summaries of GLMMs on the relationship between nectar traits and floral shape categories

**Table S11**: Summaries of GLMMs on the relationship between nectar traits and floral colour categories

**Tables S12-14**: Summaries of GLMMs corresponding to Tables S5, S10 and S11, but using total content of main sugars as a covariate, instead of sample volume.

**Table S1** List of plant species sampled, their family, flower shape and bee-colour category and type of nectar collection conducted. The washing technique collection refers to the one described by Power *et al*. (2018)

| Plant species | Family | Flower type | (Bee-) Colour category | Spectral data source | Nectar collection |
|---|---|---|---|---|---|
| ***Aconitum napellus*** | Ranunculaceae | bilabiate | uvblue | this study | microcapillary |
| ***Alchemilla conjuncta*** | Rosaceae | disk | bluegreen | this study | washing technique |
| ***Alchemilla xanthochlora*** | Rosaceae | disk | green | this study | washing technique |
| ***Androsace chamaejasme*** | Primulaceae | tube | bluegreen | this study | microcapillary |
| ***Anthyllis vulneraria*** | Fabaceae | bilabiate | green | this study | microcapillary |
| ***Aster alpinus*** | Asteraceae | head | blue | Chittka *et al.* (1994) | microcapillary |
| ***Campanula barbata*** | Campanulaceae | funnel | blue | this study | microcapillary |
| ***Campanula scheuchzeri*** | Campanulaceae | funnel | uvblue | this study | microcapillary |
| ***Carduus defloratus*** | Asteraceae | head | blue | this study | microcapillary |
| ***Carum carvi*** | Apiaceae | disk | bluegreen | this study | washing technique |
| ***Cirsium acaule*** | Asteraceae | head | blue | Chittka *et al.* (1994) | microcapillary |
| ***Clinopodium alpinum*** | Lamiaceae | bilabiate | blue | this study | microcapillary |
| ***Crepis aurea*** | Asteraceae | head | uvgreen | this study | microcapillary |
| ***Dianthus superbus*** | Caryophyllaceae | tube | blue | this study | microcapillary |
| ***Dryas octopetala*** | Rosaceae | disk | bluegreen | this study | washing technique |
| ***Echium vulgare*** | Boraginaceae | bilabiate | uvblue | this study | microcapillary |
| ***Euphrasia officinalis*** | Orobanchaceae | bilabiate | bluegreen | this study | microcapillary |
| ***Galium pumilum*** | Rubiaceae | disk | bluegreen | this study | washing technique |
| ***Globularia cordifolia*** | Plantaginaceae | head | bluegreen | this study | microcapillary |
| ***Gymnadenia nigra*** | Orchidaceae | bilabiate | uvblue | this study | microcapillary |
| ***Hippocrepis comosa*** | Fabaceae | bilabiate | green | this study | microcapillary |
| ***Hypericum maculatum*** | Hypericaceae | disk | uvgreen | this study | microcapillary |
| ***Leucanthemum vulgare*** | Asteraceae | head | bluegreen | this study | microcapillary |
| ***Lotus alpinus*** | Fabaceae | bilabiate | green | this study | microcapillary |
| ***Myosotis alpestris*** | Boraginaceae | tube | blue | this study | microcapillary |
| ***Pedicularis verticillata*** | Orobanchaceae | bilabiate | uvblue | this study | microcapillary |
| ***Phyteuma orbiculare*** | Campanulaceae | head | uvblue | this study | microcapillary |
| ***Pilosella officinarum*** | Asteraceae | head | bluegreen | this study | microcapillary |

| | | | | | |
|---|---|---|---|---|---|
| ***Prunella grandiflora*** | Lamiaceae | bilabiate | uvblue | this study | microcapillary |
| ***Ranunculus acris*** | Ranunculaceae | disk | uvgreen | this study | microcapillary |
| ***Rhododendron ferrugineum*** | Ericaceae | funnel | blue | this study | microcapillary |
| ***Salvia pratensis*** | Lamiaceae | bilabiate | uvblue | Chittka *et al.* (1994) | microcapillary |
| ***Saxifraga paniculata*** | Saxifragaceae | disk | bluegreen | this study | washing technique |
| ***Scabiosa lucida*** | Caprifoliaceae | head | blue | this study | microcapillary |
| ***Sedum album*** | Crassulaceae | disk | bluegreen | Chittka *et al.* (1994) | microcapillary |
| ***Silene vulgaris*** | Caryophyllaceae | tube | bluegreen | Chittka *et al.* (1994) | microcapillary |
| ***Thymus serpyllum*** | Lamiaceae | bilabiate | blue | this study | microcapillary |
| ***Trifolium montanum*** | Fabaceae | bilabiate | bluegreen | this study | microcapillary |
| ***Trifolium pallescens*** | Fabaceae | bilabiate | bluegreen | this study | microcapillary |
| ***Trifolium pratense*** | Fabaceae | bilabiate | bluegreen | this study | microcapillary |
| ***Vaccinium vitis-idaea*** | Ericaceae | funnel | bluegreen | this study | microcapillary |
| ***Veronica fruticans*** | Plantaginaceae | disk | uvblue | this study | microcapillary |
| ***Veronica officinalis*** | Plantaginaceae | disk | uvblue | this study | microcapillary |

**Methods S1**: Extraction and metabolomic analysis.

We conducted untargeted metabolomic analyses following the protocol from Lisec *et al*. (2006). To monitor potential contamination, a process blank (absolute ethanol) was included for every batch of 19 samples. To summarize, nectar samples and blanks were first centrifuged to remove any potential detritus and pollen, then completely dried from ethanol using a speed vacuum. For each nectar sample and blank, we added 1400 μl of methanol containing 8.46 μg/ml of ribitol and 8.46 μg/ml of L-norvaline as internal standards. They were then incubated at 70 °C for 10 minutes in a thermomixer at 950 rpm, and dried again to remove residual moisture. For derivatization, we performed methoxyamination by adding 50 μl of 15 mg/ml methoxyamine-HCl in anhydrous pyridine and incubating at 50 °C for 1.5 hours at 400 rpm. This was followed by silylation with 50 μl of MSFTA +1 % TMCS, incubated at 50 °C for one hour at 400 rpm.

To separate and analyse the compounds, we injected the samples into a gas chromatograph mass spectrometer (GC-MS, Agilent 7890B GC with installed splitter to FID and a 5977A MS) with a HP5-MS UI capillary column (30 m x 0.25 mm x 0.25 µm, Agilent, USA). Given that dominant sugars in nectar can easily overwhelm other metabolites, each sample was analysed twice: once in splitless mode to focus on detecting low-abundance metabolites, and once in split mode to prevent sugar overload (with a 20:1 split ratio). The separation programmes were the same for both runs: we used a temperature gradient from 70 to 325 °C at a ramp of 5 °C/min, and a constant helium flow rate of 0.7 ml/min.

Chromatograms resulting from the splitless mode were analysed using Mzmine (4.3.3) (Heuckeroth *et al*., 2024; Schmid *et al*., 2023). Specifically, raw GC-MS files in Agilent *.d format were converted to *.mzML format using MSconvert (ProteoWizard 3.0.24182) and imported into Mzmine. Mass detection was performed with a noise level of 150. Chromatograms were constructed using a cut-off of 4 consecutive scans, a minimum absolute peak height of 200, and an m/z tolerance of 0.5. After performing local minimum feature resolution and spectral deconvolution, we aligned the spectra using GC aligner package with an m/z tolerance of 0.5 and a retention time variance of 0.1 minutes. From the aligned spectra, we conducted a library search using the NIST14 database to identify the compounds. Duplicated features and known contaminants were excluded from further analysis, resulting in the identification of 178 entities. We manually verified 70 automated identifications to the chemical class level or with higher resolution. Quantification was based on the target ion values generated by the Mzmine pipeline, with relative concentrations calculated as ratios to the internal standard, ribitol. Note that these target-ion based values serve as proxies for comparison between different samples, but they do not allow for comparison between different compounds, unlike the FID-based quantification from our split runs. Two samples were excluded from this analysis due to machine errors.

To analyse the predominant sugars from the split runs, we used the MassHunter software package (version B.07.00, Agilent, USA). We manually identified 22 sugar-related compounds (sugars and sugar alcohols) by comparing their relative retention index and mass spectra against the NIST14 library. For sugars with available standards in our collection (such as glucose, fructose, sucrose, turanose, palatinose, galactose, myo-inositol, sorbitol and pinitol), we ran external standards to further confirm their identification. Quantification of the sugars was performed by calculating their FID areas and converting them to concentrations based on their ratio to the area of the internal standard of ribitol. We calculated sugar amounts per flower by dividing the compound quantities by the number of flowers pooled for each sample.

Finally, for all quantification data from both splitless and split runs, we further subtracted the values of the blank controls from the sample data to account for potential contamination. All blank controls exhibited similar profiles with very few compounds compared to the samples. To ensure accounting for potential contamination (i.e., carryover), we conservatively subtracted the maximum values of compounds found among blanks from all samples. We calculated compound amounts per flowers pooled for each sample. Since the non-FID data do not reflect absolute amounts, we further scaled these values from 0 to 1 by dividing each compound by its maximum value to facilitate result visualization in figures.

**Table S2** Summary table of records of visitation events to the sampled species (order as in phylogenetic tree), showing only the 35 species with 5 or more visitation records.

| Plant species | Bee visits | Non-bee hymenopteran visits | Dipteran visits | Lepidopteran visits | Coleopteran visits |
|---|---|---|---|---|---|
| *Gymnadenia nigra* | 8 | 3 | 8 | 7 | 1 |
| *Ranunculus acris* | 5 | 2 | 40 | 0 | 2 |
| *Aconitum napellus* | 11 | 2 | 3 | 0 | 1 |
| *Saxifraga paniculata* | 0 | 5 | 9 | 1 | 1 |
| *Sedum album* | 0 | 2 | 2 | 0 | 1 |
| *Hypericum maculatum* | 2 | 0 | 20 | 3 | 0 |
| *Dryas octopetala* | 0 | 0 | 6 | 0 | 1 |
| *Alchemilla xanthochlora* | 0 | 1 | 9 | 0 | 0 |
| *Alchemilla conjuncta* | 0 | 3 | 17 | 6 | 2 |
| *Hippocrepis comosa* | 20 | 1 | 3 | 1 | 0 |
| *Lotus alpinus* | 25 | 1 | 4 | 4 | 2 |
| *Anthyllis vulneraria* | 32 | 0 | 5 | 3 | 1 |
| *Trifolium pratense* | 35 | 0 | 11 | 11 | 0 |
| *Trifolium pallescens* | 12 | 0 | 1 | 0 | 0 |
| *Trifolium montanum* | 8 | 2 | 1 | 2 | 0 |
| *Dianthus superbus* | 1 | 0 | 3 | 1 | 0 |
| *Rhododendron ferrugineum* | 14 | 2 | 10 | 1 | 6 |
| *Scabiosa lucida* | 14 | 0 | 14 | 73 | 4 |
| *Carum carvi* | 1 | 3 | 14 | 0 | 7 |
| *Campanula barbata* | 15 | 0 | 6 | 0 | 7 |
| *Phyteuma orbiculare* | 5 | 1 | 8 | 15 | 1 |
| *Campanula scheuchzeri* | 18 | 0 | 4 | 7 | 8 |
| *Aster alpinus* | 0 | 0 | 14 | 12 | 1 |
| *Carduus defloratus* | 7 | 0 | 5 | 16 | 5 |
| *Leucanthemum vulgare* | 0 | 5 | 33 | 5 | 7 |
| *Pilosella officinarum* | 1 | 0 | 5 | 1 | 1 |
| *Crepis aurea* | 6 | 0 | 9 | 1 | 2 |
| *Galium pumilum* | 0 | 0 | 10 | 1 | 1 |
| *Echium vulgare* | 20 | 0 | 1 | 0 | 0 |
| *Myosotis alpestris* | 0 | 0 | 6 | 0 | 1 |
| *Veronica fruticans* | 0 | 0 | 5 | 0 | 0 |
| *Salvia pratensis* | 2 | 14 | 1 | 2 | 2 |
| *Prunella grandiflora* | 23 | 0 | 1 | 4 | 0 |
| *Clinopodium alpinum* | 12 | 0 | 2 | 3 | 0 |
| *Thymus serpyllum* | 32 | 0 | 35 | 23 | 1 |

**Methods S2**: Visual model parameters used for estimating bee-colours, using the R package “pavo 2” (version 2.9.0; Maia *et al*., 2019).

```
vismodel(spectral_data, visual = "apis",
                        achromatic = "l",
                        qcatch = "Ei",
                        illum = "D65",
                        bkg = "green",
                        vonkries = TRUE,
                        relative = FALSE)
```

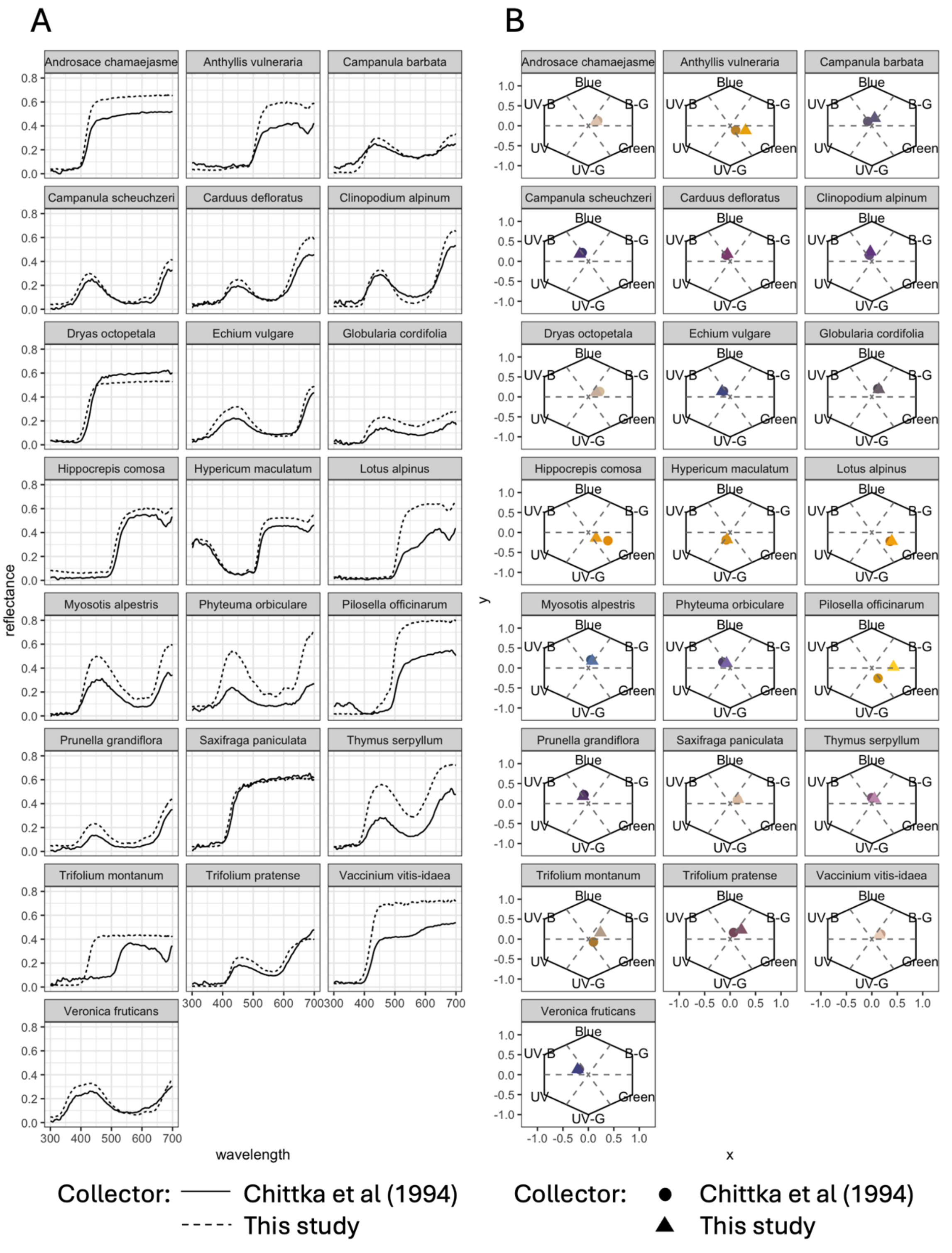


**Figure S1**: Comparison of A) spectra and B) estimated colour (in the bee-colour space) of 22 species in common between the species measured in this study and the ones reported by Chittka *et al* (1994).

**Table S3** List of metabolic components detected in nectar samples. Individual components are displayed with their putative identity and associated phylogenetic signals across the overall plant community. Compounds were identified via either solely a library search on MZmine (MZ), library search and class retention time window MZRT), library search and further confirmation with published retention indices published in the NIST14 library (MR), or by comparison to an external standard (ExS). Note that compounds classified as sugars also contain sugar derivatives. Phylogenetic signal tests on individual compounds were performed on presence/absence data via the D statistic, except for fructose, glucose and sucrose, in which presence/absence was not informative. In these cases, the phylogenetic signal was calculated via Pagel's lambda on the (log-transformed) species-average amount per flower, using the FID data.

| Method - data | Compound code | Category | Compound best match | Compound identification | Phylogenetic signal test | Phylogenetic signal | Phylogenetic signal p-value (FDR-corrected) |
|---|---|---|---|---|---|---|---|
| MS - metabolomic | X30 | Primary | Diacetone alcohol | MR | D | 1.195 | **0.000** |
| MS - metabolomic | X36 | Defensive | Phenol | MR | D | 1.903 | 0.177 |
| MS - metabolomic | X43 | Primary | Lactic acid | MR | D | 0.498 | 0.072 |
| MS - metabolomic | X47 | Primary | Lactic acid | MR | D | 1.056 | **0.004** |
| MS - metabolomic | X52 | FA | Hexanoic acid | MR | D | 1.137 | **0.000** |
| MS - metabolomic | X88 | Defensive | Oxalic acid | MR | D | 0.293 | 0.245 |
| MS - metabolomic | X102 | Primary | Dimethyl tartrate | MR | D | 1.042 | **0.000** |
| MS - metabolomic | X143 | Other | 4-methyl-2-Penten-2,4-diol | MR | D | 1.068 | **0.000** |
| MS - metabolomic | X144 | AA | Glycine | MZ | D | 0.955 | **0.004** |
| MS - metabolomic | X153 | FA | Octanoic acid | MZ | D | 0.721 | **0.049** |
| MS - metabolomic | X157 | Other | Allylamine | MZ | D | 1.157 | **0.004** |
| MS - metabolomic | X163 | Defensive | 2-acetylcyclopentanone | MZ | D | 1.035 | **0.016** |
| MS - metabolomic | X165 | Sugar | Glycerol | MR | D | 1.019 | **0.000** |
| MS - metabolomic | X191 | Sugar | Glyceric acid | MR | D | 0.914 | **0.045** |
| MS - metabolomic | X192 | AA | Proline | MZ | D | 1.092 | 0.051 |
| MS - metabolomic | X197 | FA | Nonanoic acid | MR | D | 1.086 | **0.000** |
| MS - metabolomic | X209 | Pheromone or allomone | 2-Octene-2-ol | MZ | D | 0.987 | **0.016** |
| MS - metabolomic | X223 | Phytohormone | 5-methyl-1H-Indole-3-acetic acid | MZ | D | 0.861 | **0.010** |

| | | | | | | | |
|---|---|---|---|---|---|---|---|
| MS - metabolomic | X225 | Sugar | Malic acid | MR | D | 0.839 | 0.133 |
| MS - metabolomic | X267 | Sugar | L-threitol | MR | D | 1.170 | **0.004** |
| MS - metabolomic | X278 | FA | Similar to, but not hexanoic acid | MZ | D | 1.133 | **0.004** |
| MS - metabolomic | X284 | Primary | Cadaverine | MR | D | 1.313 | **0.000** |
| MS - metabolomic | X285 | Sugar | 2,3,4-Trihydroxybutyric acid | MZ | D | 3.209 | **0.034** |
| MS - metabolomic | X286 | Sugar | Threonic acid | MZ | D | 0.250 | 0.305 |
| MS - metabolomic | X290 | AA | 2-Aminoadipic acid | MZ | D | 1.048 | **0.000** |
| MS - metabolomic | X292 | Sugar | Ribose | MR | D | 0.852 | **0.007** |
| MS - metabolomic | X297 | AA | Phenylalanine | MR | D | 1.024 | **0.000** |
| MS - metabolomic | X302 | Primary | Xylopyranose | MR | D | 0.919 | **0.000** |
| MS - metabolomic | X305 | Sugar | Arabinose | MR | D | 1.346 | **0.018** |
| MS - metabolomic | X326 | Primary | Ribonic acid | MR | D | 1.055 | **0.018** |
| MS - metabolomic | X339 | Sugar | Ribofuranose | MR | D | 0.705 | **0.015** |
| MS - metabolomic | X367 | Sugar | Meso-Erythritol | MZ | D | 0.827 | 0.059 |
| MS - metabolomic | X412 | Primary | Citric acid | MR | D | 1.067 | **0.036** |
| MS - metabolomic | X453 | Sugar | 1,5-anhydroglucitol | MZ | D | 0.681 | 0.124 |
| MS - metabolomic | X457 | Sugar | Gulose | MZ | D | 0.686 | 0.059 |
| MS - metabolomic | X466 | Defensive | Quininic acid | MR | D | 0.451 | 0.124 |
| MS - metabolomic | X472 | Sugar | Fructose | ExS | D | NA | NA |
| MS - metabolomic | X483 | Sugar | Talose | MZ | D | 1.086 | **0.000** |
| MS - metabolomic | X484 | Sugar | Glucose | ExS | D | NA | NA |
| MS - metabolomic | X494 | Sugar | Talose | MZ | D | 0.699 | 0.091 |
| MS - metabolomic | X498 | Sugar | Talose | MZ | D | 0.811 | **0.011** |
| MS - metabolomic | X519 | Sugar | Glucuronic acid lactone | MZ | D | 1.309 | **0.000** |
| MS - metabolomic | X523 | Sugar | Galactose | MZ | D | 1.149 | **0.018** |
| MS - metabolomic | X540 | Sugar | Sorbitol | MZ | D | 1.024 | **0.004** |
| MS - metabolomic | X547 | Sugar | Glucitol | MZ | D | 0.484 | 0.076 |
| MS - metabolomic | X563 | Sugar | Myo-inositol | MZ | D | 0.603 | 0.091 |
| MS - metabolomic | X568 | Sugar | Allofuranose | MZ | D | 1.188 | **0.011** |
| MS - metabolomic | X593 | Sugar | N-Acetyl-glucosamine | MZ | D | 0.433 | 0.150 |
| MS - metabolomic | X594 | Sugar | Xylofuranose | MZ | D | 0.670 | **0.018** |
| MS - metabolomic | X602 | Sugar | Pinitol | MZ | D | -0.855 | 0.808 |
| MS - metabolomic | X604 | Primary | Gluconic acid | MR | D | 1.033 | **0.000** |
| MS - metabolomic | X605 | FA | Palmitic acid | MR | D | 0.903 | **0.007** |

| | | | | | | | |
|---|---|---|---|---|---|---|---|
| MS - metabolomic | X619 | Sugar | Glucose | ExS | D | -0.735 | 0.750 |
| MS - metabolomic | X632 | Sugar | Myo-inositol | MR | D | -0.980 | 0.945 |
| MS - metabolomic | X646 | Sugar | Myo-inositol | MR | D | 0.463 | 0.177 |
| MS - metabolomic | X649 | Sugar | Myo-inositol | MZ | D | 1.060 | 0.138 |
| MS - metabolomic | X654 | Sugar | Myo-inositol | MZ | D | -0.521 | 0.920 |
| MS - metabolomic | X673 | FA | Stearic acid | MR | D | 0.819 | **0.004** |
| MS - metabolomic | X679 | FA | Hexanedioic acid, bis(2-ethylhexyl) ester | MR | D | 1.037 | **0.018** |
| MS - metabolomic | X706 | Sugar | Sucrose | ExS | D | NA | NA |
| MS - metabolomic | X729 | Sugar | Allofuranose | MZ | D | 0.786 | **0.011** |
| MS - metabolomic | X773 | Sugar | Turanose | MZ | D | 0.907 | **0.000** |
| MS - metabolomic | X849 | Sugar | Turanose | ExS | D | 0.764 | **0.016** |
| MS - metabolomic | X850 | Sugar | Palatinose | MZ | D | 0.424 | 0.150 |
| MS - metabolomic | X852 | Sugar | Mannobiose | MZ | D | 0.904 | **0.000** |
| MS - metabolomic | X910 | Sugar | Maltitol | MZ | D | 0.750 | **0.022** |
| MS - metabolomic | X913 | Sugar | Gentiobiose | MZ | D | 0.429 | 0.127 |
| MS - metabolomic | X916 | Sugar | Maltose | MZ | D | 1.037 | **0.004** |
| MS - metabolomic | X919 | Sugar | Galactinol | MZ | D | -1.426 | 1.000 |
| MS - metabolomic | X921 | Sugar | Glyceryl-glycoside | MZ | D | -0.956 | 0.804 |
| FID - sugars | pinitol | sugar alcohol | Pinitol | ExS | D | -0.093 | 0.628 |
| FID - sugars | myoinositol | sugar alcohol | Myoinositol | ExS | D | 0.626 | **0.023** |
| FID - sugars | turanose | disaccharide | Turanose | ExS | D | 0.827 | **0.011** |
| FID - sugars | sorbitol | sugar alcohol | Sorbitol | ExS | D | 0.058 | 0.503 |
| FID - sugars | palatinose | disaccharide | Palatinose | ExS | D | 1.348 | **0.036** |
| FID - sugars | disac_2 | disaccharide | Similar to, but not turanose | MZRT | D | 0.826 | **0.015** |
| FID - sugars | maltose | disaccharide | Maltose | ExS | D | 1.296 | **0.000** |
| FID - sugars | monosac_1 | monosaccharide | Allose | MZRT | D | -0.176 | 0.745 |
| FID - sugars | disac_4 | disaccharide | Similar to, but not turanose | MZRT | D | 1.115 | **0.007** |
| FID - sugars | disac_1 | disaccharide | Gentiobiose | MZRT | D | 0.878 | **0.011** |
| FID - sugars | disac_3 | disaccharide | Similar to, but not turanose | MZRT | D | 0.833 | **0.015** |
| FID - sugars | monosac_2 | monosaccharide | Allose | MZRT | D | -0.478 | 0.767 |
| FID - sugars | disac_7 | disaccharide | Maltitol | MZRT | D | 0.981 | **0.037** |

| | | | | | | | |
|---|---|---|---|---|---|---|---|
| FID - sugars | disac_5 | disaccharide | Cellobiose | MZRT | D | 1.044 | **0.016** |
| FID - sugars | disac_9 | disaccharide | Talose | MZRT | D | 0.613 | 0.390 |
| FID - sugars | alcoh_3 | sugar alcohol | Similar to, but not myo-inositol | MZRT | D | 0.498 | 0.591 |
| FID - sugars | alcoh_1 | sugar alcohol | Similar to, but not pinitol | MZRT | D | 0.436 | 0.227 |
| FID - sugars | alcoh_2 | sugar alcohol | Similar to, but not myo-inositol | MZRT | D | 0.980 | 0.177 |
| FID - sugars | disac_8 | disaccharide | Mannobiose | MZRT | D | 1.023 | **0.032** |
| FID - sugars | glucose | monosaccharide | Glucose | ExS | Pagel's lambda | 0.000 | 1.000 |
| FID - sugars | fructose | monosaccharide | Fructose | ExS | Pagel's lambda | 0.000 | 1.000 |
| FID - sugars | sucrose | disaccharide | Sucrose | ExS | Pagel's lambda | 0.000 | 1.000 |

**Table S4** Phylogenetic signal on aggregated nectar traits. Traits were averaged per species before the calculation of phylogenetic signals.

| Trait | Data transformation | Pagel's lambda | p-value |
|---|---|---|---|
| Sucrose proportion | logit | 0.000 | 1.0000 |
| Uncommon sugar richness (from FID) | log | 0.000 | 1.0000 |
| Metabolomic richness (from MS) | log | 0.000 | 1.0000 |
| Uncommon sugar dispersion (distance to median) | none | 0.014 | 0.8914 |
| Metabolomic dispersion (distance to median) | none | 0.000 | 1.0000 |
| Mean nectar volume per flower | log | 0.803 | 0.0600 |
| Mean total sugars per flower | log | 0.000 | 1.0000 |

**Table S5**. Generalised Linear Mixed Models (GLMMs) testing for correlation between visitation PCs and the 5 aggregated nectar traits. CI refers to Confidence Interval (95%), and PG refers to Pairwise Groups based on post-hoc Tukey comparisons ($p < 0.05$). Models account for site and sample nectar volume as fixed covariates, with plant species specified random effect levels. * In the case of analysis resulting from splitless data, 2 samples were excluded due to machine errors.

| **Response variable:** | **Sucrose proportion** | | | **Metabolomic richness** | | | **Uncommon sugar richness** | | | **Metabolomic dispersion** | | | **Uncommon sugar dispersion** | | |
|---|---|---|---|---|---|---|---|---|---|---|---|---|---|---|---|
| *Predictors* | *Estimate* ***(odds** ratio)* | *CI* | *p-value* | *Estimate* ***(log-link)*** | *CI* | *p-value* | *Estimate* ***(log-link)*** | *CI* | *p-value* | *Estimate* ***(log-link)*** | *CI* | *p-value* | *Estimate* ***(log-link)*** | *CI* | *p-value* |
| (Intercept) | 0.56 | 0.35 – 0.90 | **0.016** | 2.42 | 2.35 – 2.49 | **<0.001** | 0.74 | 0.54 – 0.93 | **<0.001** | -0.81 | -0.86 – -0.76 | **<0.001** | -1.57 | -1.77 – -1.36 | **<0.001** |
| PC1 | 1.71 | 1.24 – 2.36 | **0.001** | 0.06 | 0.02 – 0.10 | **0.005** | 0.20 | 0.07 – 0.33 | **0.002** | -0.02 | -0.05 – 0.01 | 0.116 | 0.15 | 0.02 – 0.29 | **0.027** |
| PC2 | 1.24 | 0.82 – 1.87 | 0.318 | 0.02 | -0.03 – 0.07 | 0.439 | 0.09 | -0.07 – 0.24 | 0.258 | 0.02 | -0.02 – 0.05 | 0.33 | 0.08 | -0.09 – 0.25 | 0.348 |
| PC3 | 0.9 | 0.56 – 1.45 | 0.675 | -0.06 | -0.12 – -0.00 | **0.044** | -0.16 | -0.34 – 0.03 | 0.099 | 0.02 | -0.02 – 0.06 | 0.398 | -0.12 | -0.31 – 0.08 | 0.248 |
| site [2000] | 0.72 | 0.60 – 0.87 | **0.001** | -0.07 | -0.13 – 0.00 | 0.054 | -0.01 | -0.11 – 0.09 | 0.809 | -0.07 | -0.12 – -0.02 | **0.008** | -0.07 | -0.17 – 0.02 | 0.135 |
| vol nectar (uL) | | | | 0.06 | 0.03 – 0.09 | **<0.001** | 0.14 | 0.11 – 0.17 | **<0.001** | 0.05 | 0.03 – 0.08 | **<0.001** | 0.08 | 0.05 – 0.12 | **<0.001** |
| **Random Effects** | | | | | | | | | | | | | | | |
| $\sigma^2$ | 0.17 | | | 6.92 | | | 1.13 | | | 0.00 | | | 0.01 | | |
| $\tau_{00}$ | 1.79 $_{species}$ | | | 0.02 $_{species}$ | | | 0.22 $_{species}$ | | | 0.01 $_{species}$ | | | 0.33 $_{species}$ | | |
| ICC | 0.68 | | | 0.00 | | | 0.16 | | | 0.65 | | | 0.97 | | |
| N | 35 $_{species}$ | | | 35 $_{species}$ | | | 35 $_{species}$ | | | 35 $_{species}$ | | | 35 $_{species}$ | | |
| Observations | 250 | | | 248 * | | | 250 | | | 248 * | | | 250 | | |
| Marginal $R^2$ / Conditional $R^2$ | 0.249 / 0.934 | | | 0.003 / 0.005 | | | 0.097 / 0.243 | | | 0.302 / 0.755 | | | 0.233 / 0.979 | | |

**Table S6.** Distance-based Redundancy Analysis (dbRDA) partitioning the variance of nectar metabolomic profiles across plant species with available visitation data (n = 35). The analysis is based on a species-level Jaccard distance matrix derived from metabolite presence/absence. The model evaluates the marginal effects of pollinator visitation (PC1) and maximum nectar volume analysed per species, while conditioning out the effects of phylogeny via principal components (PhyloPC1–PC5, accounting for 92.66 % of variation in phylogenetic distances). Adjusted $R^2$ represents the total variance explained by the unconditioned predictors. p-values for unconstrained predictors are derived from 999 marginal permutations, while the significance of the conditioned phylogenetic structure is evaluated via partial model comparison against a phylogeny-free baseline. Significant effects (p < 0.05) are highlighted in bold. * The effect of the conditioned phylogeny was evaluated via partial model comparison against a phylogeny-free baseline model.

| **Response variable** | **Metabolomic (Jaccard) distance matrix** | | | |
|---|---|---|---|---|
| **Formula** | distance_matrix ~ visitation_PC1 + maximum_nectar_volume_per_species + condition(PhyloPC1 + PhyloPC2 + PhyloPC3 + PhyloPC4 + PhyloPC5) | | | |
| *Predictors* | *df* | *Sum of squares* | *F* | *p-value* |
| Conditioned phylogeny (PhyloPC1-PC5) * | 5 | 1.22 | 1.28 | **0.001** |
| Visitation PC1 | 1 | 0.43 | 2.26 | **0.002** |
| Maximum nectar volume per species | 1 | 0.19 | 0.99 | 0.459 |
| Unconstrained residuals | 27 | 5.18 | | |
| Observations | 35 | | | |
| R2 adjusted | 0.083 | | | |

**Table S7**: Vector fitting results identifying 22 individual nectar metabolites correlated with the dbRDA ordination (p-value < 0.05; see model in **Table S6 and Figure 4A** in main text). For compound names see **Table S3**. Vectors were fitted using the envfit function (vegan package; Oksanen *et al*., 2025). The coordinates (dbRDA1 and dbRDA2) represent the directions of the fitted vectors, indicating their alignment with the ordination axes (visitation PC1 and maximum nectar volume analysed per species). The squared correlation coefficient (r2) quantifies the strength of the association for each compound to the ordinated space. P-values are derived from 999 permutations of the data, and metabolites are organized by ascending p-value.

| **Metabolite code (presence)** | **dbRDA1** | **dbRDA2** | **r2** | ***p-value*** |
|---|---|---|---|---|
| **X483** | 0.83518 | -0.54998 | 0.3445 | **0.001** |
| **X523** | 0.81874 | -0.57416 | 0.3776 | **0.001** |
| **X594** | 0.83758 | 0.54632 | 0.4574 | **0.001** |
| **X673** | 0.90365 | 0.42827 | 0.4072 | **0.001** |
| **X729** | 0.99567 | -0.09293 | 0.6473 | **0.001** |
| **X850** | 0.93777 | 0.34725 | 0.4121 | **0.001** |
| **X852** | 0.7071 | 0.70712 | 0.3098 | **0.001** |
| **X910** | 0.95268 | -0.30398 | 0.426 | **0.001** |
| **X913** | 0.9934 | -0.11467 | 0.623 | **0.001** |
| **X916** | 0.95766 | -0.28791 | 0.5486 | **0.001** |
| **X197** | 0.12629 | 0.99199 | 0.371 | **0.002** |
| **X302** | 0.99963 | 0.0272 | 0.3764 | **0.002** |
| **X165** | 0.56498 | 0.82511 | 0.2542 | **0.006** |
| **X278** | 0.49145 | 0.8709 | 0.2568 | **0.008** |
| **X773** | 0.9335 | -0.35858 | 0.2383 | **0.014** |
| **X646** | 0.99927 | 0.03814 | 0.2259 | **0.022** |
| **X163** | 0.61519 | 0.78838 | 0.2007 | **0.027** |
| **X679** | 0.61519 | 0.78838 | 0.2007 | **0.027** |
| **X604** | 0.98045 | 0.19676 | 0.1983 | **0.030** |
| **X292** | 0.99942 | 0.03412 | 0.202 | **0.032** |
| **X849** | 0.99859 | -0.05313 | 0.1937 | **0.040** |
| **X547** | 0.90869 | 0.41748 | 0.1734 | **0.049** |

**Table S8**: Summary of GLMM testing for association between visitation PC1 and floral shape categories**.**

| | **PC 1** | | | |
|---|---|---|---|---|
| *Predictors* | *Estimates* | *CI* | *p-value* | *Pairwise group* |
| (Intercept)<br>flower type [bilabiate] | 1.44 | 1.05 – 1.83 | **<0.001** | **a** |
| flower type [tube] | -2.40 | -3.48 – -1.33 | **<0.001** | bc |
| flower type [head] | -1.55 | -2.22 – -0.89 | **<0.001** | **b** |
| flower type [funnel] | -1.37 | -2.27 – -0.46 | **0.004** | **b** |
| flower type [disk] | -3.07 | -3.66 – -2.47 | **<0.001** | **c** |
| Observations | 35 | | | |
| $R^2$ / $R^2$ adjusted | 0.794 / 0.767 | | | |

**Table S9**: Summary of GLMM testing for association between visitation PC1 and floral colour categories**.**

| | **PC 1** | | | |
|---|---|---|---|---|
| *Predictors* | *Estimates* | *CI* | *p-value* | *Pairwise group* |
| (Intercept)<br>section [blue] | 0.01 | -2.06 – 2.08 | 0.993 | **a** |
| section [bluegreen] | -1.72 | -3.01 – -0.44 | **0.010** | **b** |
| section [green] | -1.18 | -3.07 – 0.71 | 0.213 | ab |
| section [uvblue] | 0.95 | -0.11 – 2.02 | 0.078 | **a** |
| section [uvgreen] | -1.1 | -2.68 – 0.47 | 0.162 | ab |
| Observations | 35 | | | |
| $R^2$ / $R^2$ adjusted | 0.348 / 0.261 | | | |

**Table S10:** GLMM outputs evaluating the association of floral shape categories with the five aggregated nectar traits. The intercept level corresponds to bilabiate flowers. CI: Confidence Interval; PG: Pairwise Groups based on post-hoc Tukey comparisons ($p < 0.05$). * In the case of analysis resulting from splitless data, 2 samples were excluded due to machine errors.

**Response variable: Sucrose proportion**

Model formula: variable ~ flower shape + site + (1|species)

| *Predictors* | *Estimate* ***(odds** ratio)* | *CI* | *p-value* | *PG* |
|---|---|---|---|---|
| flower type [bilabiate; intercept] | 1.01 | 0.46 – 2.23 | 0.985 | n.s. |
| flower type [tube] | 0.11 | 0.02 – 0.62 | **0.012** | n.s. |
| flower type [head] | 0.35 | 0.10 – 1.26 | 0.108 | n.s. |
| flower type [funnel] | 0.80 | 0.14 – 4.53 | 0.805 | n.s. |
| flower type [disk] | 0.24 | 0.07 – 0.83 | **0.023** | n.s. |
| site [2000] | 0.73 | 0.61 – 0.87 | **<0.001** | |
| vol nectar (uL) | | | | |
| $\sigma^2$ | 1.66 | | | |
| $\tau_{00}$ | 2.36 species | | | |
| ICC | 0.77 | | | |
| N | 43 species | | | |
| Observations | 285 | | | |
| Marginal $R^2$ / Conditional $R^2$ | 0.179 / 0.946 | | | |

**Response variable: Metabolomic richness**

Model formula: variable ~ flower shape + site + nectar_volume_uL+ (1|species)

| *Predictors* | *Estimate* ***(log-link)*** | *CI* | *p-value* | *PG* |
|---|---|---|---|---|
| flower type [bilabiate; intercept] | 2.48 | 2.40 – 2.57 | **<0.001** | **a** |
| flower type [tube] | -0.02 | -0.19 – 0.15 | 0.854 | ab |
| flower type [head] | 0.05 | -0.07 – 0.16 | 0.433 | **a** |
| flower type [funnel] | 0.12 | -0.04 – 0.27 | 0.137 | **a** |
| flower type [disk] | -0.23 | -0.35 – -0.11 | **<0.001** | **b** |
| site [2000] | -0.09 | -0.15 – -0.02 | **0.007** | |
| vol nectar (uL) | 0.06 | 0.03 – 0.09 | **<0.001** | |
| $\sigma^2$ | 6.52 | | | |
| $\tau_{00}$ | 0.01 species | | | |
| ICC | 0.00 | | | |
| N | 43 species | | | |
| Observations | 283 * | | | |
| Marginal $R^2$ / Conditional $R^2$ | 0.003 / 0.005 | | | |

**Response variable: Uncommon sugar richness**

Model formula: variable ~ flower shape + site + nectar_volume_uL+ (1|species)

| *Predictors* | *Estimate* ***(log-link)*** | *CI* | *p-value* | *PG* |
|---|---|---|---|---|
| flower type [bilabiate; intercept] | 0.98 | 0.72 – 1.24 | **<0.001** | **a** |
| flower type [tube] | -0.57 | -1.18 – 0.05 | 0.07 | ab |
| flower type [head] | -0.30 | -0.72 – 0.13 | 0.17 | ab |
| flower type [funnel] | -0.04 | -0.58 – 0.51 | 0.897 | ab |
| flower type [disk] | -0.6 | -1.02 – -0.19 | **0.005** | **b** |
| site [2000] | -0.02 | -0.12 – 0.07 | 0.66 | |
| vol nectar (uL) | 0.14 | 0.11 – 0.17 | **<0.001** | |
| $\sigma^2$ | 1.03 | | | |
| $\tau_{00}$ | 0.21 species | | | |
| ICC | 0.17 | | | |
| N | 43 species | | | |
| Observations | 285 | | | |
| Marginal $R^2$ / Conditional $R^2$ | 0.075 / 0.233 | | | |

**Response variable: Metabolomic dispersion**

Model formula: variable ~ flower shape + site + nectar_volume_uL+ (1|species)

| *Predictors* | *Estimate* ***(log-link)*** | *CI* | *p-value* | *PG* |
|---|---|---|---|---|
| flower type [bilabiate; intercept] | -0.85 | -0.92 – -0.78 | **<0.001** | n.s. |
| flower type [tube] | 0.07 | -0.07 – 0.21 | 0.339 | n.s. |
| flower type [head] | 0.00 | -0.10 – 0.11 | 0.973 | n.s. |
| flower type [funnel] | 0.06 | -0.08 – 0.20 | 0.399 | n.s. |
| flower type [disk] | 0.02 | -0.08 – 0.12 | 0.732 | n.s. |
| site [2000] | -0.06 | -0.11 – -0.01 | **0.02** | |
| vol nectar (uL) | 0.05 | 0.03 – 0.08 | **<0.001** | |
| $\sigma^2$ | 0.00 | | | |
| $\tau_{00}$ | 0.01 species | | | |
| ICC | 0.70 | | | |
| N | 43 species | | | |
| Observations | 283 * | | | |
| Marginal $R^2$ / Conditional $R^2$ | 0.209 / 0.759 | | | |

**Response variable: Uncommon sugar dispersion**

Model formula: variable ~ flower shape + site + nectar_volume_uL+ (1|species)

| *Predictors* | *Estimate* ***(log-link)*** | *CI* | *p-value* | *PG* |
|---|---|---|---|---|
| flower type [bilabiate; intercept] | -1.36 | -1.62 – -1.09 | **<0.001** | n.s. |
| flower type [tube] | -0.40 | -1.01 – 0.20 | 0.194 | n.s. |
| flower type [head] | -0.19 | -0.62 – 0.24 | 0.39 | n.s. |
| flower type [funnel] | -0.02 | -0.59 – 0.55 | 0.935 | n.s. |
| flower type [disk] | -0.56 | -0.98 – -0.13 | **0.010** | n.s. |
| site [2000] | -0.09 | -0.18 – 0.01 | 0.085 | |
| vol nectar (uL) | 0.09 | 0.05 – 0.12 | **<0.001** | |
| $\sigma^2$ | 0.01 | | | |
| $\tau_{00}$ | 0.23 species | | | |
| ICC | 0.97 | | | |
| N | 43 species | | | |
| Observations | 285 | | | |
| Marginal $R^2$ / Conditional $R^2$ | 0.218 / 0.975 | | | |

**Table S11:** GLMM outputs evaluating the association of floral bee-colour categories with the five aggregated nectar traits. The intercept level corresponds to green flowers. CI: Confidence Interval; PG: Pairwise Groups based on post-hoc Tukey comparisons ($p < 0.05$). * In the case of analysis resulting from splitless data, 2 samples were excluded due to machine errors.

| **Response variable:** | **Sucrose proportion** | | | | **Metabolomic richness** | | | | **Uncommon sugar richness** | | | | **Metabolomic dispersion** | | | | **Uncommon sugar dispersion** | | | |
|---|---|---|---|---|---|---|---|---|---|---|---|---|---|---|---|---|---|---|---|---|
| Model formula: | variable ~ flower colour + site + (1\|species) | | | | variable ~ flower colour + site + nectar_volume_uL+ (1\|species) | | | | variable ~ flower colour + site + nectar_volume_uL+ (1\|species) | | | | variable ~ flower colour + site + nectar_volume_uL+ (1\|species) | | | | variable ~ flower colour + site + nectar_volume_uL+ (1\|species) | | | |
| *Predictors* | *Estimate* ***(odds** ratio)* | *CI* | *p-value* | *PG* | *Estimate* ***(log-link)*** | *CI* | *p-value* | *PG* | *Estimate* ***(log-link)*** | *CI* | *p-value* | *PG* | *Estimate* ***(log-link)*** | *CI* | *p-value* | *PG* | *Estimate* ***(log-link)*** | *CI* | *p-value* | *PG* |
| section [green; intercept] | 0.77 | 0.15 – 3.82 | 0.746 | n.s. | 2.47 | 2.31 – 2.62 | **<0.001** | ab | 0.68 | 0.18 – 1.18 | **0.008** | ab | -0.86 | -0.98 – -0.73 | **<0.001** | n.s. | -1.84 | -2.37 – -1.30 | **<0.001** | n.s. |
| section [uvblue] | 0.66 | 0.10 – 4.38 | 0.663 | n.s. | 0.04 | -0.14 – 0.22 | 0.638 | ab | 0.40 | -0.18 – 0.97 | 0.179 | **a** | 0.00 | -0.15 – 0.14 | 0.996 | n.s. | 0.60 | -0.01 – 1.21 | 0.055 | n.s. |
| section [blue] | 1.17 | 0.18 – 7.80 | 0.87 | n.s. | 0.07 | -0.10 – 0.25 | 0.427 | **a** | 0.24 | -0.34 – 0.82 | 0.419 | **a** | 0.01 | -0.13 – 0.16 | 0.839 | n.s. | 0.49 | -0.12 – 1.10 | 0.119 | n.s. |
| section [uvgreen] | 0.60 | 0.05 – 7.00 | 0.681 | n.s. | -0.06 | -0.30 – 0.17 | 0.599 | ab | -0.19 | -0.96 – 0.58 | 0.631 | ab | 0.08 | -0.11 – 0.26 | 0.406 | n.s. | -0.12 | -0.93 – 0.69 | 0.772 | n.s. |
| section [bluegreen] | 0.31 | 0.05 – 1.86 | 0.200 | n.s. | -0.14 | -0.31 – 0.03 | 0.11 | **b** | -0.26 | -0.82 – 0.30 | 0.367 | **b** | 0.04 | -0.10 – 0.17 | 0.602 | n.s. | 0.06 | -0.53 – 0.64 | 0.851 | n.s. |
| site [2000] | 0.72 | 0.60 – 0.87 | **<0.001** | | -0.07 | -0.13 – -0.01 | **0.033** | | -0.02 | -0.11 – 0.07 | 0.678 | | -0.05 | -0.10 – -0.00 | **0.037** | | -0.08 | -0.18 – 0.01 | 0.089 | |
| vol nectar (uL) | | | | | 0.06 | 0.03 – 0.09 | **<0.001** | | 0.14 | 0.11 – 0.17 | **<0.001** | | 0.06 | 0.03 – 0.08 | **<0.001** | | 0.08 | 0.05 – 0.12 | **<0.001** | |
| $\sigma^2$ | 1.66 | | | | 6.55 | | | | 1.02 | | | | 0.00 | | | | 0.01 | | | |
| $\tau_{00}$ | 2.61 $_{species}$ | | | | 0.02 $_{species}$ | | | | 0.20 $_{species}$ | | | | 0.01 $_{species}$ | | | | 0.22 $_{species}$ | | | |
| ICC | 0.84 | | | | 0.00 | | | | 0.16 | | | | 0.69 | | | | 0.97 | | | |
| N | 43 $_{species}$ | | | | 43 $_{species}$ | | | | 43 $_{species}$ | | | | 43 $_{species}$ | | | | 43 $_{species}$ | | | |
| Observations | 285 | | | | 283 * | | | | 285 | | | | 283 * | | | | 285 | | | |
| Marginal $R^2$ / Conditional $R^2$ | 0.102 / 0.946 | | | | 0.002 / 0.005 | | | | 0.087 / 0.236 | | | | 0.206 / 0.757 | | | | 0.296 / 0.975 | | | |

**Table S12**. Corresponding model outputs to Table S5 (GLMM outputs on visitation PCs vs. aggregated nectar traits), but with the per-sample total amount of the three main sugars (sucrose, fructose, glucose; FID-quantified) used as a covariate in place of sample nectar volume, to further confirm that instrumental detection limits did not drive the associations found for richness and dispersion metrics. Sucrose proportion tests were excluded because this trait should not be influenced by instrumental detection limits. Response variables, model structure, random effects, and abbreviations follow Table S5

| | **Metabolomic richness** | | | **Uncommon sugar richness** | | | **Metabolomic dispersion** | | | **Uncommon sugar dispersion** | | |
|---|---|---|---|---|---|---|---|---|---|---|---|---|
| *Predictors* | *Estimate (log-link)* | *CI* | *p-value* | *Estimate (log-link)* | *CI* | *p-value* | *Estimate (log-link)* | *CI* | *p-value* | *Estimate (log-link)* | *CI* | *p-value* |
| (Intercept) | 2.42 | 2.35 – 2.49 | **<0.001** | 0.74 | 0.55 – 0.93 | **<0.001** | -0.81 | -0.86 – -0.75 | **<0.001** | -1.58 | -1.78 – -1.37 | **<0.001** |
| PC1 | 0.06 | 0.02 – 0.10 | **0.003** | 0.20 | 0.07 – 0.32 | **0.002** | -0.02 | -0.05 – 0.01 | 0.213 | 0.15 | 0.01 – 0.28 | **0.031** |
| PC2 | 0.03 | -0.02 – 0.07 | 0.303 | 0.10 | -0.05 – 0.26 | 0.195 | 0.02 | -0.01 – 0.06 | 0.217 | 0.09 | -0.08 – 0.25 | 0.320 |
| PC3 | -0.05 | -0.11 – 0.00 | 0.067 | -0.14 | -0.33 – 0.04 | 0.129 | 0.02 | -0.02 – 0.06 | 0.331 | -0.11 | -0.30 – 0.09 | 0.281 |
| total sugars (µg) | 0.31 | 0.11 – 0.50 | **0.002** | 0.83 | 0.64 – 1.02 | **<0.001** | 0.27 | 0.12 – 0.43 | **0.001** | 0.56 | 0.35 – 0.78 | **<0.001** |
| site [2000] | -0.07 | -0.14 – 0.00 | 0.053 | -0.05 | -0.15 – 0.04 | 0.296 | -0.07 | -0.12 – -0.02 | **0.005** | -0.09 | -0.18 – 0.00 | 0.063 |
| $\sigma^2$ | 7.03 | | | 1.11 | | | 0.00 | | | 0.01 | | |
| $\tau_{00}$ | 0.02 species | | | 0.22 species | | | 0.01 species | | | 0.34 species | | |
| ICC | 0.00 | | | 0.16 | | | 0.71 | | | 0.97 | | |
| N | 35 species | | | 35 species | | | 35 species | | | 35 species | | |
| Observations | 248 | | | 250 | | | 248 | | | 250 | | |
| Marginal $R^2$ / Conditional $R^2$ | 0.002 / 0.005 | | | 0.099 / 0.245 | | | 0.246 / 0.778 | | | 0.232 / 0.980 | | |

**Table S13.** Corresponding model outputs to Table S10 (floral shape categories vs. aggregated nectar traits), with per-sample total main-sugar amount as covariate instead of nectar volume. All other specifications follow Table S10.

| **Response variable:** | **Metabolomic richness** | | | | **Uncommon sugar richness** | | | | **Metabolomic dispersion** | | | | **Uncommon sugar dispersion** | | | |
|---|---|---|---|---|---|---|---|---|---|---|---|---|---|---|---|---|
| Model formula: | variable ~ flower shape + site + nectar total sugars + (1\|species) | | | | variable ~ flower shape + site + nectar total sugars + (1\|species) | | | | variable ~ flower shape + site + nectar total sugars + (1\|species) | | | | variable ~ flower shape + site + nectar total sugars + (1\|species) | | | |
| *Predictors* | *Estimate* ***(log-link)*** | *CI* | *p-value* | *PG* | *Estimate* ***(log-link)*** | *CI* | *p-value* | *PG* | *Estimate* ***(log-link)*** | *CI* | *p-value* | *PG* | *Estimate* ***(log-link)*** | *CI* | *p-value* | *PG* |
| flower type [bilabiate; intercept] | 2.48 | 2.40 – 2.56 | **<0.001** | **a** | 0.98 | 0.72 – 1.23 | **<0.001** | **a** | -0.85 | -0.92 – -0.77 | **<0.001** | n.s. | -1.38 | -1.65 – -1.11 | **<0.001** | n.s. |
| flower type [tube] | -0.01 | -0.18 – 0.16 | 0.939 | ab | -0.53 | -1.14 – 0.07 | 0.084 | ab | 0.07 | -0.08 – 0.22 | 0.334 | n.s. | -0.37 | -0.98 – 0.23 | 0.228 | n.s. |
| flower type [head] | 0.05 | -0.07 – 0.17 | 0.416 | **a** | -0.28 | -0.70 – 0.13 | 0.184 | ab | 0.00 | -0.11 – 0.11 | 0.996 | n.s. | -0.18 | -0.61 – 0.26 | 0.426 | n.s. |
| flower type [funnel] | 0.14 | -0.02 – 0.29 | 0.082 | **a** | 0.04 | -0.50 – 0.57 | 0.892 | ab | 0.08 | -0.07 – 0.22 | 0.295 | n.s. | 0.03 | -0.55 – 0.60 | 0.931 | n.s. |
| flower type [disk] | -0.23 | -0.35 – -0.11 | **<0.001** | **b** | -0.60 | -1.01 – -0.19 | **0.004** | **b** | 0.01 | -0.09 – 0.12 | 0.836 | n.s. | -0.54 | -0.97 – -0.12 | **0.012** | n.s. |
| site [2000] | -0.09 | -0.15 – -0.03 | **0.006** | | -0.06 | -0.15 – 0.03 | 0.186 | | -0.07 | -0.11 – -0.02 | **0.01** | | -0.10 | -0.20 – -0.01 | **0.036** | |
| total sugars (µg) | 0.34 | 0.16 – 0.52 | **<0.001** | | 0.84 | 0.66 – 1.02 | **<0.001** | | 0.28 | 0.12 – 0.43 | **<0.001** | | 0.58 | 0.37 – 0.80 | **<0.001** | |
| $\sigma^2$ | 6.62 | | | | 1.01 | | | | 0 | | | | 0.01 | | | |
| $\tau_{00}$ | 0.01 species | | | | 0.20 species | | | | 0.01 species | | | | 0.23 species | | | |
| ICC | 0.00 | | | | 0.17 | | | | 0.71 | | | | 0.97 | | | |
| N | 43 species | | | | 43 species | | | | 43 species | | | | 43 species | | | |
| Observations | 283 | | | | 285 | | | | 283 | | | | 285 | | | |
| Marginal $R^2$ / Conditional $R^2$ | 0.003 / 0.004 | | | | 0.077 / 0.233 | | | | 0.169 / 0.763 | | | | 0.224 / 0.976 | | | |

**Table S14.** Corresponding model outputs to Table S11 (floral shape categories vs. aggregated nectar traits), with per-sample total main-sugar amount as covariate instead of nectar volume. All other specifications follow Table S11.

| **Response variable:** | **Metabolomic richness** | | | | **Uncommon sugar richness** | | | | **Metabolomic dispersion** | | | | **Uncommon sugar dispersion** | | | |
|---|---|---|---|---|---|---|---|---|---|---|---|---|---|---|---|---|
| Model formula: | variable ~ flower colour + site + nectar total sugars + (1\|species) | | | | variable ~ flower colour + site + nectar total sugars + (1\|species) | | | | variable ~ flower colour + site + nectar total sugars + (1\|species) | | | | variable ~ flower colour + site + nectar total sugars + (1\|species) | | | |
| *Predictors* | *Estimate **(log-link)*** | *CI* | *p-value* | *PG* | *Estimate **(log-link)*** | *CI* | *p-value* | *PG* | *Estimate **(log-link)*** | *CI* | *p-value* | *PG* | *Estimate **(log-link)*** | *CI* | *p-value* | *PG* |
| section [green; intercept] | 2.47 | 2.31 – 2.62 | **<0.001** | ab | 0.70 | 0.21 – 1.20 | **0.005** | ab | -0.86 | -0.99 – -0.72 | **<0.001** | n.s. | -1.84 | -2.38 – -1.30 | **<0.001** | n.s. |
| section [uvblue] | 0.04 | -0.14 – 0.22 | 0.66 | ab | 0.36 | -0.21 – 0.93 | 0.219 | **a** | 0.00 | -0.15 – 0.15 | 0.997 | n.s. | 0.57 | -0.04 – 1.18 | 0.069 | n.s. |
| section [blue] | 0.08 | -0.10 – 0.25 | 0.395 | **a** | 0.24 | -0.33 – 0.81 | 0.407 | ab | 0.02 | -0.13 – 0.17 | 0.776 | n.s. | 0.49 | -0.12 – 1.10 | 0.116 | n.s. |
| section [uvgreen] | -0.06 | -0.31 – 0.03 | 0.628 | ab | -0.19 | -0.96 – 0.57 | 0.616 | ab | 0.08 | -0.11 – 0.28 | 0.412 | n.s. | -0.11 | -0.92 – 0.70 | 0.784 | n.s. |
| section [bluegreen] | -0.14 | -0.31 – 0.01 | 0.098 | **b** | -0.28 | -0.83 – 0.27 | 0.320 | **b** | 0.03 | -0.11 – 0.17 | 0.675 | n.s. | 0.05 | -0.54 – 0.64 | 0.867 | n.s. |
| site [2000] | -0.07 | -0.13 – -0.01 | **0.030** | | -0.06 | -0.15 – 0.03 | 0.203 | | -0.06 | -0.11 – 0.01 | **0.020** | | -0.10 | -0.19 – -0.00 | **0.039** | |
| total sugars (µg) | 0.31 | 0.12 – 0.51 | **0.001** | | 0.83 | 0.65 – 1.01 | **<0.001** | | 0.28 | 0.13 – 0.44 | **0.001** | | 0.57 | 0.35 – 0.79 | **<0.001** | |
| $\sigma^2$ | 6.65 | | | | 1.00 | | | | 0 | | | | 0.01 | | | |
| $\tau_{00}$ | 0.02 $_{species}$ | | | | 0.20 $_{species}$ | | | | 0.01 $_{species}$ | | | | 0.22 $_{species}$ | | | |
| ICC | 0.00 | | | | 0.16 | | | | 0.72 | | | | 0.97 | | | |
| N | 43 $_{species}$ | | | | 43 $_{species}$ | | | | 43 $_{species}$ | | | | 43 $_{species}$ | | | |
| Observations | 283 | | | | 285 | | | | 283 | | | | 285 | | | |
| Marginal $R^2$ / Conditional $R^2$ | 0.002 / 0.005 | | | | 0.090 / 0.238 | | | | 0.160 / 0.762 | | | | 0.301 / 0.977 | | | |

**References**


Chittka, L., Shmida, A., Troje, N., and Menzel, R. 1994. Ultraviolet as a component of flower reflections, and the colour perception of hymenoptera. *Vision research* 34:1489–1508.

Heuckeroth, S., Damiani, T., Smirnov, A., Mokshyna, O., Brungs, C., Korf, A., Smith, J. D., Stincone, P., Dreolin, N., Nothias, L.-F., *et al*. (2024). Reproducible mass spectrometry data processing and compound annotation in mzmine 3. *Nature protocols* 19:2597–2641.

Lisec, J., Schauer, N., Kopka, J., Willmitzer, L., and Fernie, A. R. (2006). Gas chromatography mass spectrometry–based metabolite profiling in plants. *Nature protocols* 1:387–396.

Maia, R., Gruson, H., Endler, J. A., and White, T. E. 2019. pavo 2: new tools for the spectral and spatial analysis of colour in r. *Methods in Ecology and Evolution* 10.

Oksanen, J., Simpson, G. L., Blanchet, F. G., Kindt, R., Legendre, P., Minchin, P. R., *et al.*, 2025. vegan: Community Ecology Package. URL https://vegandevs.github.io/vegan/, r601 package version 2.7-0, https://github.com/vegandevs/vegan.

Power, E. F., Stabler, D., Borland, A. M., Barnes, J., and Wright, G. A. 2018. Analysis of nectar from low-volume flowers: A comparison of collection methods for free amino acids. *Methods in ecology and evolution* 9:734–743.

Schmid, R., Heuckeroth, S., Korf, A., Smirnov, A., Myers, O., Dyrlund, T. S., *et al*. (2023). Integrative analysis of multimodal mass spectrometry data in mzmine 3. *Nature biotechnology* 41:447–449.
